# Nuclear alpha-synuclein is present in the human brain and is modified in dementia with Lewy bodies

**DOI:** 10.1101/2021.10.20.465125

**Authors:** David J. Koss, Daniel Erskine, Andrew Porter, Pawel Palmoski, Marta Leite, Johannes Attems, Tiago F. Outeiro

## Abstract

Dementia with Lewy bodies is pathologically defined by the cytoplasmic accumulation of alpha-synuclein within neuronal cells in the brain. Alpha-synuclein is predominately pre-synaptic, but has been reported present in various subcellular compartments in cell and animal models. In particular, nuclear alpha-synuclein is evident *in-vitro* and in disease models and has been associated with altered DNA integrity, gene transcription, nuclear homeostasis. However, owing to various factors, the presence of alpha-synuclein in the nuclei of human brain cells remains controversial, as does its role in synucleinopathies.

Here, we close this gap and provide a unique demonstration confirming the presence of nuclear alpha-synuclein in post-mortem brain tissue obtained from cases of dementia with Lewy bodies as well as from controls via immunohistochemistry, immunoblot, and label-free mass-spectrometry.

Discrete intra-nuclear alpha-synuclein puncta reactive against phosphorylated serine 129-alpha-synuclein and pan-alpha-synuclein antibodies were observed in cortical neurons and non-neuronal cells in fixed brain sections and in isolated nuclear preparations from Dementia with Lewy bodies cases and matched controls. Subsequent biochemical analysis of subcellular fractionated tissue confirmed alpha-synuclein as present in a nuclear fraction at levels ~ 10-fold lower than in the cytoplasm. Critically, however, an increase in monomeric nuclear alpha-synuclein phosphorylated as serine 129 was observed in cases of dementia with Lewy bodies alongside higher molecular weight pan- and phosphorylation reactive alpha-synuclein species, consistent with the formation of intranuclear phosphorylated alpha-synuclein oligomers. Furthermore, the presence of nuclear alpha-synuclein was confirmed via label free mass spectrometry, as 6 unique alpha-synuclein derived peptide sequences were identified in nuclear fractions (71.4% sequence coverage).

Collectively, our data confirm the presence of nuclear alpha-synuclein in human brain tissue and describe nuclear pathology associated with dementia with Lewy bodies. These findings address a major controversy in the synucleinopathy field by confirming the presence of nuclear alpha-synuclein in autoptic human brain tissue and, for the first time, identify that alpha-synuclein is aggregated into novel and potentially pathological assemblies in the nucleus as part of the disease process associated with dementia with Lewy bodies and thus may contribute to the disease phenotype.

## Introduction

The hallmark pathology of Lewy body diseases, including Parkinson’s disease and dementia with Lewy bodies, is the intraneuronal accumulation of alpha-synuclein (aSyn) in protein inclusions known as Lewy bodies and Lewy neurites. Although Lewy pathology is prominently localised within brain stem, limbic, or cortical regions and is consistent with the symptomology of the diseases, inconsistencies between aggregate burden, toxicity, cell type-specific vulnerability and overall disease severity are well documented.^1–6^ Therefore, whether Lewy pathology (Lewy bodies and Lewy neurites) is the underlying factor for neuronal dysfunction and death seen in Lewy body diseases is still unclear. Nevertheless, genetic association studies linking mutations in the gene encoding for aSyn (*SNCA*) with Parkinson’s Disease^7^ and an array of studies in cellular and animal models, support the hypothesis that dysfunction of aSyn might play a causative role in Lewy body diseases.^8,9^

aSyn is generally assumed to be simply a presynaptic protein. Although initial studies had indicated that the protein may also be present within the nucleus,^10^ this has long been highly controversial and overlooked.^11^ Indeed reports of prominent nuclear aSyn (aSyn^Nuc^) have been undermined by the discovery that several key commercial aSyn antibodies possess cross-reactivity with a putative non-aSyn nuclear antigen.^11^ However, increased support for the nuclear localisation of aSyn has been recently gained by studies utilising novel antibodies validated in aSyn knockout mouse tissue, fluorophore-coupled fusion proteins, and the use of nuclear isolation protocols both in expression systems and at endogenous levels in the brain.^12–15^ Likewise although there is a lack of evidence supporting physiological aSyn^Nuc^ in humans, intranuclear aSyn inclusions are a pathological feature of multiple system atrophy and as such supports the detrimental potential of altered aSyn^Nuc^ composition in human neurodegenerative diseases.^16^

Evidence based on studies using *in vitro* and *in vivo* models, suggests that aSyn^Nuc^ regulates a number of nuclear functions, interacting directly with DNA^13,14,17^ and histones,^18^ in turn regulating gene transcription^13,14,19^ and also DNA repair.^17^ Critically, in models of disease, overexpression of wild type or disease relevant aSyn mutants (e.g. A30P, A53T, or G51D) results in enhanced levels of aSyn^Nuc^ and in the formation of intranuclear inclusions, alongside impairments in gene transcription and in nuclear import processes.^20–24^

Although nuclear dysfunction may play a major part in the pathology of human synucleinopathies, the occurrence of aSyn^Nuc^ in human brain material is still controversial. However, establishing the intra-nuclear presence of aSyn must be considered essential not only to progress our understanding of the molecular mechanisms associated with Lewy body diseases, but also to inform on the normal function of aSyn, both of which are instrumental for the design of future therapeutic strategies.

## Materials and Methods

### Tissue preparation

Post-mortem human brain tissue from clinic-pathologically confirmed cases of dementia with Lewy Bodies (n=14) and non-neurodegenerative diseases controls (n=17) for comparison was obtained from the Newcastle Brain Tissue Resource. For in-situ tissue histology, cingulate or lateral temporal cortex slide-mounted sections were prepared from the fixed right hemisphere of the brain (4% paraformaldehyde immersion fixed, 6-week). Frozen temporal grey matter (Broadman area 21/22, middle and superior temporal gyrus, 500 mg) for subcellular fractionation, biochemical analysis, and mass spectrometry was obtained from the contralateral left hemisphere of the brain, which was snap frozen between copper plates at - 120°C, before sample extraction.

Conformation of disease was established for each case based on a review of clinical history following death and pathological assessment of brain tissue according to Lewy Body Braak staging^25^ and McKeith Criteria^26,27^ as well the National institute of ageing – Alzheimer’s Association (NIA-AA) criteria,^28^ including neurofibrillary tangle Braak staging,^29^ Thal Aß phases^30^ and Consortium to Establish a Registry for Alzheimer’s Disease (CERAD) scoring.^31^ For details of cohort and tissue use see Table 1 and Supplementary Table 1.

**Table 1.** Human tissue cohort. Human cases use for histology, western blot and mass spectrometry. Cases are separated by disease classification according to non-diseased controls (Con) and dementia with Lewy bodies (DLB). Case numbers (n), sex, age, postmortem interval (PMI), neurofibrillary tangle (NFT) Braak stage, Thal phase, Consortium to Establish a Registry for Alzheimer’s Disease (CERAD), the National Institute of Ageing – Alzheimer’s Association (NIA-AA) criteria, Lewy body (LB) Braak stage and McKeith criteria are provided. For age and PMI both range and mean ±SEM are provided. For numerical scores of pathology, range and percentage composition are given. For CERAD scores, negative (neg), A and B reported. For NIA-AA, not, low and intermediate (inter) risk for Alzheimer’s disease. For McKeith criteria, only percentage composition is given, where cases free of Lewy bodies (No LB) and neocortical predominate (Neo) are indicated. *=based on available data.

| Disease | n | Sex (% male) | Age (years) | PMI (hrs) | NFT Braak stage | Thal Phase | CERAD | NIA-AA | LB Braak stage | McKeith Criteria |
| --- | --- | --- | --- | --- | --- | --- | --- | --- | --- | --- |
| Histology |  |  |  |  |  |  |  |  |  |  |
| Con | 6 | 66.7% | 60-90<br>82±4.5 | 24-85<br>55.3±9.5 | 0-IV<br>16.6%-0<br>16.6%-I<br>33.3%-II<br>16.6%-III<br>16.6%-IV | 0-3<br>16.6%-0<br>16.6%-1<br>16.6%-2<br>50%-3 | neg-B<br>83.3%-neg<br>16.6%-B | Low-inter<br>83.3%-low<br>16.6%-inter | 0<br>100%-0 | 100%-No LB |
| DLB | 9 | 66.7% | 72-91<br>78.2±1.9 | 10-99<br>55±12.1 | I-III<br>11.1%-I<br>89.9%-III | 0-4<br>14.3%-0<br>14.3%-1<br>14.3%-2<br>14.3%-3<br>42.9%-4 | *neg-B<br>55.6%-neg<br>22.2%-A<br>22.2%-B | not-inter<br>11.1%-not<br>55.6%-low<br>33.3%-inter | *6<br>100%-6 | 100%-Neo |
| Western blot |  |  |  |  |  |  |  |  |  |  |
| Con | 9 | 55.6% | 73-99<br>85.3±2.7 | 5-75<br>31.7±7.4 | 0-III<br>22.2%-0<br>11.1%-I<br>55.6%-II<br>11.1%-III | 0-4<br>22.2%-0<br>22.2%-1<br>11.1%-2<br>33.3%-3<br>11.1%-4 | neg<br>100%-neg | not-low<br>22.2%-not<br>77.7%-low | 0<br>100%-0 | 100%-No LB |
| DLB | 8 | 75% | 68-92<br>78.5±3 | 8-99<br>39.5±11.9 | II-III<br>37.5%-II<br>62.5%-III | 3-5<br>12.5%-3<br>62.5%-4<br>25%-5 | neg-B<br>62.5%-neg<br>25%-A<br>12.5%-B | low-inter<br>87.5%-low<br>12.5%-inter | 5-6<br>25%-5<br>75%-6 | 100%-Neo |
| Mass spectrometry |  |  |  |  |  |  |  |  |  |  |
| Con | 3 | 33.3% | 64-99<br>81±10.1 | 5-93<br>41±26. | I-II<br>33.3%-I<br>66.7%-II | 0-1<br>66.7%-0<br>33.3%-1 | neg<br>100%-neg | not-low<br>66.7%-low<br>33.3%-inter | 0<br>100%-0 | 100%-No LB |
| DLB | 3 | 100% | 78-81<br>79.7±0.9 | 8-34<br>22.7±7.7 | III<br>100%-III | 3-4<br>33.3%-3<br>67.7%-4 | A-B<br>33.3%-A<br>67.7%-B | Low-inter<br>33.3%-low<br>66.7%-inter | 6<br>100%-6 | 100%-Neo |

Forebrain tissue samples from three month-old aSyn knockout mice, generated as previously outlined,^32^ were obtained, snap frozen in liquid nitrogen, and stored at −80°C prior to use.

### Histochemical detection of nuclear aSyn

Paraffin embedded slide-mounted tissue sections (6-10 μm thick) were dewaxed (5 mins, xylene emersion), rehydrated (5 mins, descending concentrations of ethanol; 99, 95, 75%, emersion) and washed in Tris-buffered saline (5 mM Tris, 145mM NaCl, pH 7.4). In order to determine optimal conditions for the detection of nuclear antigens, prior to antibody staining, sections were subject to a variety of antigen retrieval methods (Citrate buffer, Formic acid, EDTA and /or Proteinase K treatment, see Table 2 for full details). Sections were subsequently blocked in 5% normal goat serum containing Tris-buffered saline (1hr, RT), labelled for phospho-Ser129 aSyn (mouse pS129 IgG2,^33^ 1:1000 or EP1536Y [ab51253, Abcam], 1:500 dilution), pan aSyn (N-terminal directed, SYN303 [aa 2-12 Cat# 82401, Biolegend], 1:500, non-amyloid β component directed Syn-1 [aa91-99, Cat# 610787, BD Trasduction Laboratories], 1:500), Histone H3 ([Cat# 3638S, Cell Signalling], 1:500) and/or NeuN ([ab104224, Abcam], 1:200) via an overnight incubation in primary antibody containing solution (5% normal goat serum containing Tris-buffered saline, 4°C) followed by secondary antibody incubation (1hr, RT, goat anti-rabbit alexa 594 and goat anti-mouse alexa 488, 1:500, Fisher Scientific). Stained sections were treated with Sudan Black (0.03% Sudan Black B in 70% ethanol) to quench endogenous fluorescence and coverslipped with DAPI containing Prolong Diamond Mountant (Fisher Scientific). Antibody labelled sections were imaged via a wide-field fluorescence microscope system (Nikon Eclipse 90i microscope, DsQi1Mc camera and NIS elements software V 3.0, Nikon) or via confocal microscope (Lecia SP 8, LAS X software, Leica-microsystems).

**Table 2.** Antigen retrieval methods. Protocols for each of the antigen retrieval methods evaluated for nuclear antigen detection.

| Antigen retrieval | Method |
| --- | --- |
| Citrate | Submersion in pre-heated citrate buffer (10 mM Citric acid, 0.05 % Tween 20, pH 6) with microwave heat-assisted antigen retrieval (800W, 10 mins). |
| Formic | Submersion in 90% Formic acid for 1 hr at room temperature (RT). |
| Proteinase K | Slides incubated at 37°C for 20 mins in Proteinase K (0.6u/ml) containing TE buffer (50 mM Tris, 1mM EDTA, 0.5% Triton X-100, pH 8). |
| EDTA + Formic | Submersion in pre-heated EDTA buffer (10mM EDTA, pH 8) and subsequent heating in pressure cooker (heated under maximum pressure for 2 mins) followed by submersion in 90% formic acid for 10 mins at RT. |

### Tissue homogenisation and fractionation

Purified nuclear extracts were generated as previously described.^34^ Tissue blocks (250 mg) and individual cerebral hemispheres from aSyn knockout mice were lysed 1:16 (W:V) in nuclear extraction buffer (in mM; 0.32 sucrose, 5 CaCl_2_, 3 Mg(Ac)_2_ 10 Tris-HCl, 0.1% NP-40, pH 8, containing cOmplete protease inhibitor cocktail and phostop tablets, 1/10ml, Sigma) via dounce homogenisation to generate crude whole tissue lysates. Cytoplasmic fractions were obtained from the centrifugation (800 rcf x 40mins, 4°C) of crude lysates (500 μl) and retention of the resulting supernatant. Nuclei were isolated from the remaining lysates via sucrose gradient centrifugation (1.8 M Sucrose, 3mM Mg(Ac)_2_, 10 mM Tris-HCl, pH 8, 107,000 rcf x 2.5hrs, 4°C). Pelleted nuclei were washed to remove any potential cytoplasmic contamination and resuspended in 500 μl 0.01M phosphate buffer saline. Generated fractions were either directly applied to microscope slides for immunochemical staining (as outlined above) or frozen at −80°C until use.

### Immunoblotting analyses

Nuclear fractions, pre-treated with DNAse (50u/ml, 15min RT, RQ1 RNAase free DNAse, Promega) and cytoplasmic fractions were adjusted, according to Bradford assay (Sigma) to 80 ng/μl with LDS sample buffer (Fisher Scientific), reducing agent (Fisher Scientific) and dH_2_O and heated at 70°C for 10mins. Denatured, reduced samples (4 μg/lane for phospho-aSyn blots and 1.2 μg/lane for all other blots) were separated via SDS page electrophoresis (200 V, 35 mins in MES buffer for aSyn detection) and transferred to nitrocellulose membranes (0.2 μm) via Iblot (7 mins, 20 mV; Fisher Scientific). Initial blots were tested for optimisations of aSyn retention as labelled by MJFR1 either by pre-blocking heat treatment (microwaved, 800W, 5 mins in boiling 10mM phosphate buffered saline) or via 30 mins chemical fixation (4% paraformaldehyde, 30mins, RT). All subsequent blots labelled for aSyn were first pre-treated via microwave-heat assisted protein retention (microwaved, 800W, 5 mins in boiling 10 mM phosphate buffered saline), prior to being blocked in 5% milk powder containing Tris buffered saline with Tween-20 (5 mM Tris, 145 mM NaCl, 0.01% Tween-20, pH 7.4, 1hr, RT). Membranes were stained for aSyn (pS129-aSyn, EP1536Y, 1:1000, Syn-1, 1:5000 and C-terminal directed MJRF1 [aa119-123, Cat# ab138501, Abcam], 1:5000) in Tris buffered saline with Tween-20 supplemented with 0.05% Sodium Azide overnight at 4°C and labelled with secondary antibodies (goat anti-mouse HRP or goat anti-rabbit-HRP, 1:5000, in 5% milk powder containing Tris buffered saline with Tween-20, 1hr, RT). Between each incubation step, blots were washed 3x 5 min in Tris buffered saline with Tween-20). Antigen-antibody interactions were visualised via enhanced chemiluminescence (1.25 mM Luminol, 30 μM coumaric acid and 0.015% H_2_O_2_), images captured with a Fuji LAS 4000 with imaging software (Fuji LAS Image, Raytek, Sheffield, UK) and saved in 16 bit for analysis and 8 bit for illustration. Protein loading and fractionation purity was establishing via reprobing membranes with the cytoplasmic marker glyceraldehyde 3-phosphate dehydrogenase (GAPDH; 1:5,000, 14C10, Cell signalling) and the nuclear marker Histone H3 (1:5000).

### Mass spectrometry analyses of fractionated tissue

Frozen samples (250 mg) were fractionated as above, with minor amendments, such that following nuclear isolation, pellets were resuspended in 50μl of 5% SDS containing phosphate buffered saline and cytoplasmic fractions were mixed 1:1 with 10% SDS solution (Thermofisher) to give a final 5% SDS concentration. Protein digestion was carried out using S-Trap (Protifi). Samples were heated at 95°C and sonicated to remove DNA/ RNA. They were then reduced with tris(2-carboxyethyl)phosphine (5mM final concentration, 15min incubation at 55 °C) and alkylated with Iodoacetamide (10mM final concentration, 10min incubation at RT). Each sample was then acidified with 12% Phosphoric acid (final concentration of 2.5%, v/v), followed by addition of 6 vol. of loading buffer (90% methanol, 100mM Tetraethylammonium bromide, pH 8) and loaded onto S-Trap cartridges. The loaded cartridges were spun at 4000xg for 30s and washed three times with 90% loading buffer and the flow through being discarded. Retained proteins were digested with trypsin (Worthington), at a ratio of 10:1 protein to trypsin, in digestion buffer (50mM Tetraethylammonium bromide, pH8.5) for 3h at 47°C.

Peptides were released of the cartridge with three sequential washes; first with 50 μl of 50mM Tetraethylammonium bromide, followed by 50μl of 0.2% formic acid and finally with 50μl of 50% acetonitrile and 0.2% formic acid. The solution was frozen then dried in a centrifugal concentrator to a volume of ~1μl. The peptide sample was reconstituted in 0.2% formic acid to an estimated concentration of 500ng/μl.

The equivalent of 500ng of the peptide sample was loaded per LCMS run, peptides were separated with a 125 min nonlinear gradient (3-40% B, 0.1% formic acid (Line A) and 80% acetonitrile, 0.1% formic acid (LineB)) using an UltiMate 3000 RSLCnano high-performance liquid chromatography. Samples were first loaded/desalted onto a 300μm x 5mm C18 PepMap C18 trap cartridge in 0.1% formic acid at 10 μl/min for 3 min and then further separated on a 75μmx50cm C18 column (Thermo EasySpray -C18 2 μm) with integrated emitter at 400nl/min. The eluent was directed to an Thermo Orbitrap Exploris 480 mass spectrometer through the EasySpray source at a temperature of 320°C, spray voltage 1900 V. The total LCMS run time was 150 min. Orbitrap full scan resolution was 60,000, RF lens 50%, Normalised ACG Target 300%. Precursors for MSMS were selected via a top 20 method. MIPS set to peptide, Intensity threshold 5.0 e3, charge state 2-7 and dynamic exclusion after 1 times for 35 s 10ppm mass tolerance. ddMS2 scans were performed at 15000 resolution, HCD collision energy 27%, first mass 110 m/z, ACG Target Standard.

The acquired DDA data (pooled from 3 controls and 3 cases of dementia with Lewy bodies, see Table 1) was searched against the human protein sequence database, available from (https://www.uniprot.org/uniprot/?query=proteome:UP000005640), concatenated to the Common Repository for Adventitious Proteins v.2012.01.01 (cRAP, ftp://ftp.thegpm.org/fasta/cRAP), using MaxQuant v1.6.43. Parameters used: cysteine alkylation: iodoacetamide, digestion enzyme: trypsin, Parent Mass Error of 5ppm, fragment mass error of 10ppm. The confidence cut-off representative to FDR<0.01 was applied to the search result file. Nuclear and cytoplasmic enrichments were verified by means of submitting the lists of identified proteins from each of the fractions to Shiny Go^35^ (V0.66).

### Statistical analysis

Immunoreactivity of western blots were quantified via area under the curve of isolated bands (ImageJ, NIH) adjusted to the area under curve of corresponding loading controls. Samples were processed in batches such that for each marker, each blot contained ≥3 control and cases of dementia with Lewy bodies, allowing the normalisation of measurements to control values within blot prior to pooling between blots. Comparisons were subsequently made via nonparametric Mann-Whitney tests, p<0.05 was deemed as significant.

Analysis of the LC-MS data (exported form MaxQuant) was performed in Perseus.^36^ First, all the common contaminants and reversed database hits were removed from the dataset, as well as proteins quantified with less than 2 unique peptides. Intensity values for each protein were transformed to log2 and the technical replicates were averaged. Median was then subtracted within each sample to account for unequal loading and the width of the distribution was adjusted. Next, the dataset was filtered, only keeping proteins which have at least 2 valid values in at least one experimental group (4 groups in total). Remaining missing values were imputed from the left tail of the normal distribution (2StDev away from the mean, +/−0.3 StDev).

Principle component analysis was conducted on the resulting dataset and Heat maps were generated (preceded by Z-score transformation, for protein across cases and fractions). Analysis of cellular components as per Shiny Go were performed with p<0.05 following FDR adjustments and the top 10 and 30 hits based on number of proteins within with a given category taken.

### Data availability

All data sets are available from the corresponding author upon request.

## Results

### In situ detection of aSyn^Nuc^

Firstly, we optimized immunofluorescence staining protocols on fixed cortical sections for the detection of nuclear antigens. Using Histone H3 as a general nuclear marker, a variety of antigen retrieval methods (see Table 2) were tested for optimal detection in post-mortem fixed human brain tissue. In comparison to individual treatment in citrate buffer, formic acid, or without pre-treatment, an antigen retrieval protocol combining pressurised heating in EDTA and 10 min submersion in 90% formic acid was found to yield the most robust H3 nuclear labelling which co-localised with dye-based DNA staining via DAPI (Supplementary Fig. 1). Consistent with the improved access of antibodies into the nuclear compartment, the nuclear labelling of pS129 with mouse anti-pS129 was greatly enhanced following a EDTA + formic acid pre-treatment, with only modest labelling of the nuclear compartment evident when either citrate buffer, proteinase K or formic acid was used alone (Fig. 1A-B). Nevertheless, under each antigen retrieval protocol, pS129 staining was localised to the nucleus within puncta at discrete sites, albeit with reduced fluorescence intensity when compared to pathological Lewy bodies and Lewy neurites. Under optimal conditions with EDTA + formic acid pre-treatment nuclear pS129 staining was evident in all nuclei and was apparent in both tissue from control and cases of dementia with Lewy bodies (Fig. 1C). Similar staining was obtained when using the commercially available pS129 (EP1536Y), such that intranuclear puncta were clearly evident in NeuN positive neurons and NeuN negative non-neuronal cells of both brain sections from controls and cases of dementia with Lewy bodies (Fig. 2A). The intranuclear localisation of this immunoreactivity signal was confirmed via confocal microscopy, orthogonal projections of z-stack images demonstrating ps129 aSyn positive puncta within DAPI positive nuclear structures (Fig. 2B).

**Figure 1.**
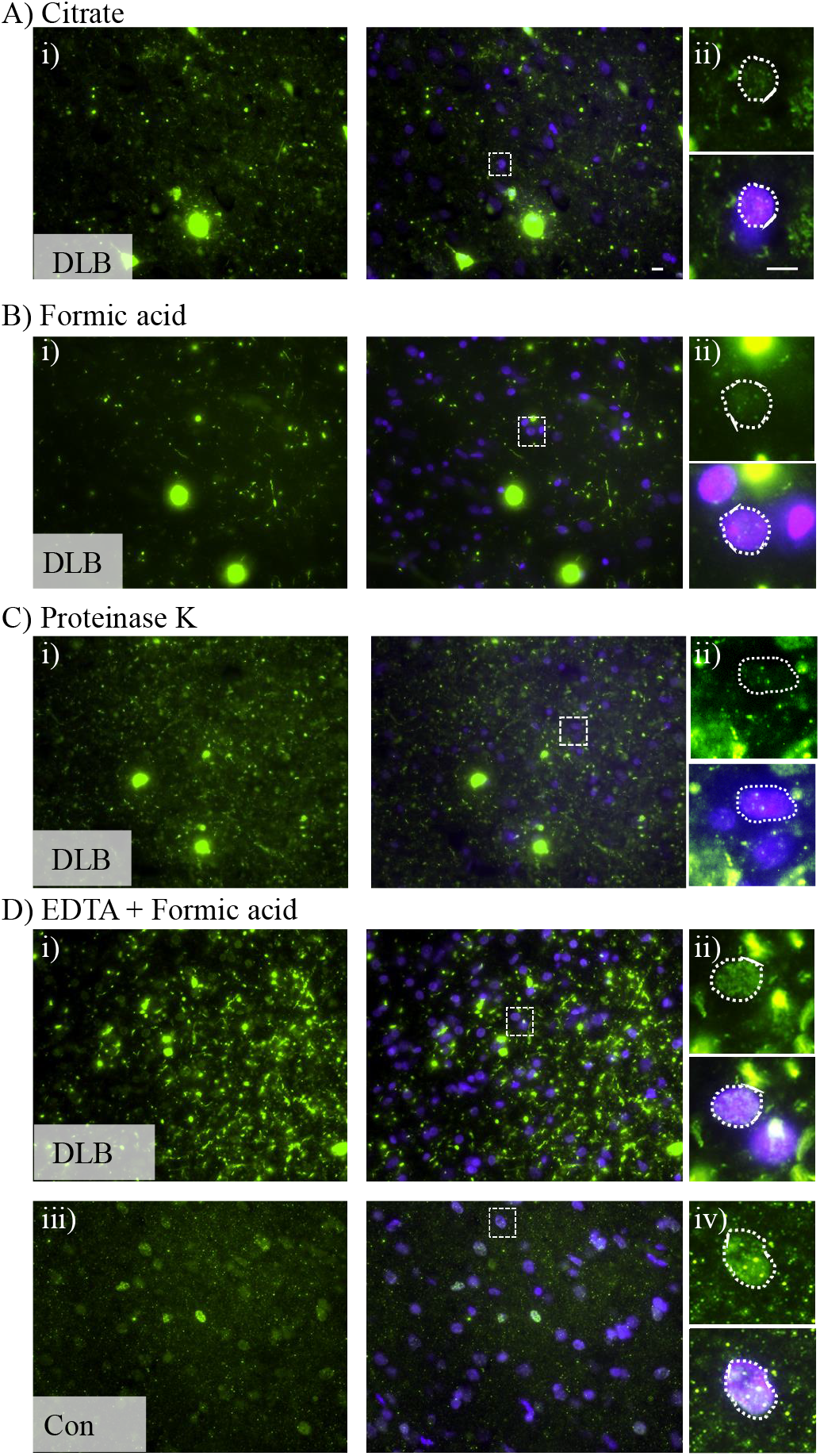
Antigen retrieval methods determine the detection of nuclear aSyn. Example micrograph images (i, x40 objective) of phospho-serine 129 positive aSyn (pS129) immunoreactivity and DAPI nuclear stain in fixed post-mortem cingulate tissue sections from cases of dementia with Lewy bodies (DLB). Sections were pre-treated prior to antibody staining with Citrate buffer (**A**), Formic acid (**B**), Protienase K (**C**) and EDTA + Formic acid (**D**) methods of antigen retrieval. Expanded area inserts (ii, dotted line boxes in i) are shown for both pS129 alone and in combination with DAPI nuclear stain, where the nuclear outline is highlighted (dotted outline). Note robust detection of punctate intranuclear staining of pS129 following EDTA + Formic acid-based antigen retrieval. In **D**, similar treated sections from a non-diseased control case (con) is also shown (iii + iv). Scale bar in i + iii = 10 μm and ii, iv =5 μm.

**Figure 2.**
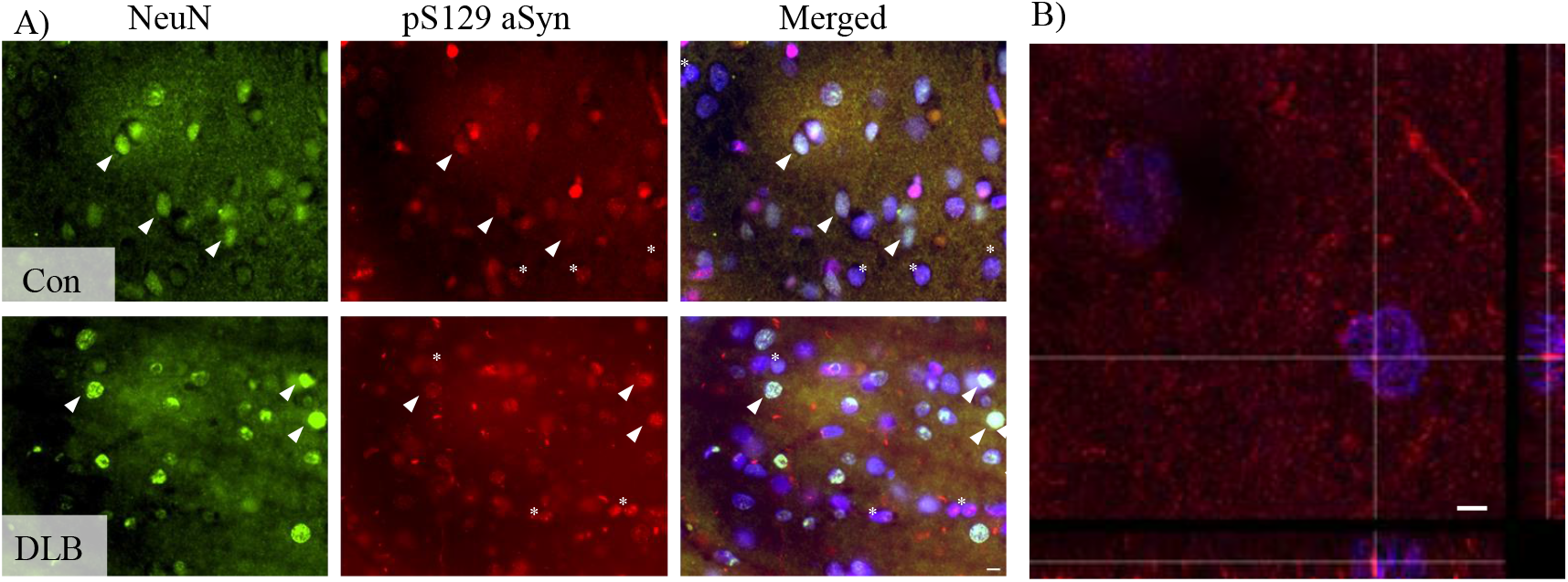
Nuclear staining phosphorylated alpha-synuclein in neuronal and nonneuronal cells. Representative images from control and dementia with Lewy bodies (DLB) cases stained with NeuN and commercial phospho-serine129-aSyn antibody (EP1536Y) following EDTA + Formic acid antigen retrieval (**A**). Note the clear punctate intranuclear staining of pS129 in NeuN positive (arrowheads) and negative cells (asterisks) evident in Con and DLB cases. Nuclear localisation of pS129 puncta was confirmed via confocal z-stack images (v, x63 objective, 3 x digital zoom) with orthogonal reconstruction, pS129 immunoreactivity localised within DAPI positive nuclei (**B**). Scale bar in a = 10 μm and in b= 5 μm.

We further confirmed genuine in-situ aSyn^Nuc^ via the labelling of additional sections with N-terminal (Syn303) and the non-amyloid component (Syn-1) directed antibodies (Fig. 3). Despite the prominent staining of the neuropil via pan-aSyn antibodies, consistent with the predominate pre-synaptic localization of the protein, numerous examples of positive aSyn immunoreactivity overlapping with the DAPI nuclear stain were evident in sections from controls and cases of dementia with Lewy bodies, without overt disease dependent changes. Again orthogonal projections demonstrated multiple aSyn positive puncta within DAPI positive nuclei (Fig. 3Biii).

**Figure 3.**
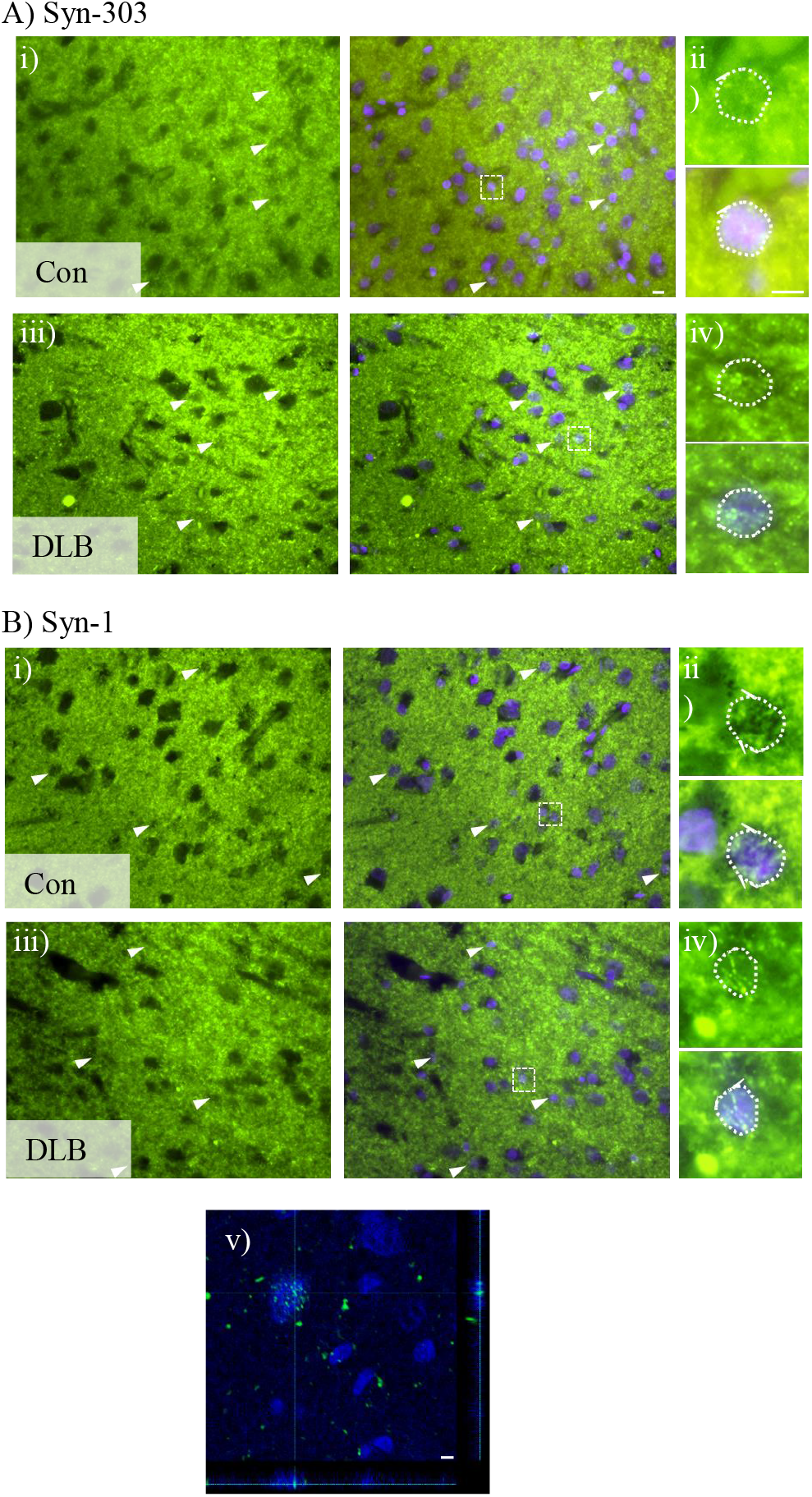
Pan-aSyn antibodies label nuclear aSyn in-situ. Example micrograph images (i and iii, x40 objective) of N-terminal directed Syn 303 antibody (**A**) and non-amyloid component directed Syn-1 (**B**) immunoreactivity and DAPI nuclear stain in cingulate human brain tissue from control (Con) and dementia with Lewy bodies cases (DLB). Sections were pre-treated prior to antibody staining with EDTA + Formic acid antigen retrieval. Expanded area inserts (ii and iv, dotted line boxes in i and iii) are shown for both Syn antibodies alone and in combination with DAPI nuclear stain, (dotted line outlines nuclei). For Syn-1, additional confocal z-stack image (iv, x20 objective with 6x digital zoom) with orthogonal reconstruction is shown, confirming intranuclear localisation of pS129 immunoreactivity. Scale bar in i + iii = 10 μm and ii, iv + v =5 μm.

### Detection of aSyn within isolated nuclei

In a bid to simplify the sample complexity, frozen temporal cortex tissue samples were used for the isolation of nuclei via sucrose gradient centrifugation. Isolated nuclear preparations were immunostained with either pS129 (mouse ps129) or pan-Syn-1 antibodies and the nuclear localisation of aSyn was confirmed via confocal microscopy (Fig. 4A). In both cases, we observed punctate immunoreactivity within the nuclei, in agreement with discrete intranuclear regions of aSyn enrichment as detected in fixed tissue histology.

**Figure 4.**
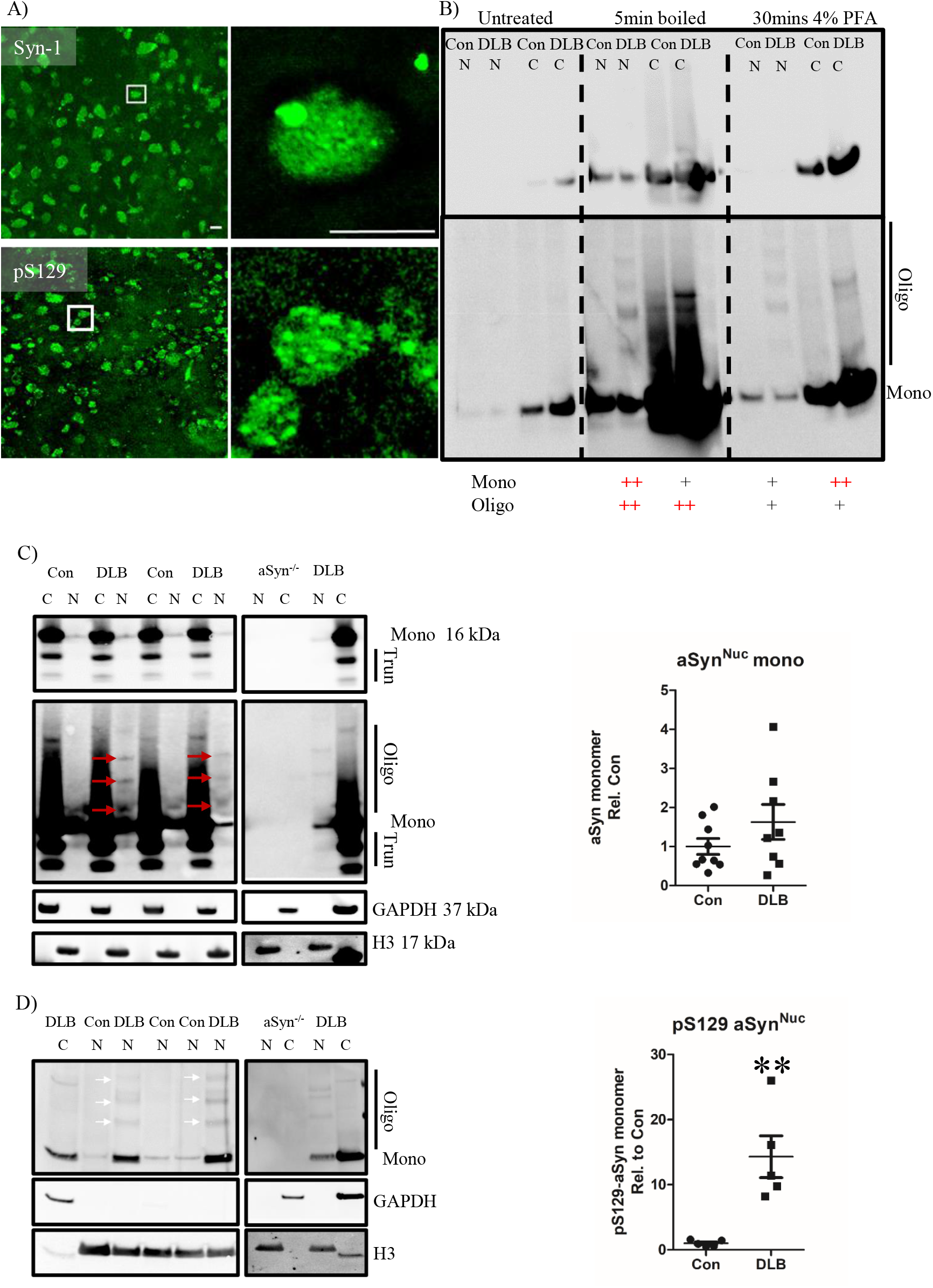
Quantification of nuclear aSyn pathology in cases of dementia with lewy bodies in isolated nuclei. **A)** Representative confocal micrograph images (x40 objective, inserts ii + iv taken with 10x digital zoom, scale = 5 μm) stained for pan-aSyn (Syn-1, i +ii) and phospho-serine 129 aSyn (pS129, iii+iv). **B)** Optimisation of immunoblot detection of aSyn from cytoplasmic (c) and nuclear (n) fractionates generated from control (Con) and dementia with Lewy bodies (DLB) cases. Comparisons of post-transfer membrane phosphate buffered saline heating (5 mins boiling) or 4% paraformaldehyde chemical (30mins 4% paraformaldehyde) fixation with non-fixed (untreated) controls is shown, captured under optimal and overexposed settings. Monomeric (mono) and oligomeric (oligo) immunoreactivity is highlighted alongside a ranking of fixation methods based retention of the aSyn species (+ = enhanced and ++ = very enhanced). Quantification of **C)** pan-aSyn via Syn-1 and **D)** pS129 via EP1536Y, (i) example immunoblots following boiling fixation are shown at optimised and overexposed capture settings and mono, oligo and truncated (trunc) species are highlighted (i). Loading controls of GAPDH and Histone H3 demonstrate fractionation purity and resulting immunoreactivity from similarly fractionated aSyn knockout mouse brain tissue (aSyn^-/-^) demonstrated antibody specificity. Quantification of nuclear aSyn monomers between DLB and controls cases is also shown (ii), demonstrating increased pS129 Syn in DLB cases compared to controls (n=5 per group), despite no change in total aSyn levels (n=9 and 8 for con and DLB respectively). Data shown as scatter plots with mean ± SEM, statistical analysis was performed via two-tailed non-parametric Mann-Whitney test, **=p=0.0033, n=5 cases per group.

The nuclear localization of aSyn was further examined via western blot analyses of isolated nuclei. Due to the relatively low protein yield of the nuclear preparations as compared to a crude cytoplasmic fractions, we first optimised the membrane retention of aSyn to enable adequate protein detection.^37^ Post-transfer membrane boiling in phosphate buffered saline and chemical fixation with 4% paraformaldehyde enhanced aSyn immunoreactivity in both nuclear and cytoplasmic tissue fractions (Fig. 4B). In particular, enhancement of the aSyn signal was further increased after boiling in phosphate buffered saline. Furthermore, the pan-aSyn (MJFR1) antibody detected higher molecular weight aSyn species only in the nuclear fractions isolated from cases of dementia with Lewy bodies, and not in the same fractions from controls (Fig. 4B). Notably the two membrane fixation methods favoured the retention of different aSyn species, such that paraformaldehyde treatment was optimal for cytoplasmic aSyn monomers, whist phosphate buffered saline treatment was optimal for nuclear monomers and higher molecular weight species (Fig. 4B), likely suggesting differing chemical composition between the different pools of aSyn.

The nuclear presence of aSyn was further confirmed in a larger number of cases, following the boiling of membranes in phosphate buffered saline and aSyn detection via pan-Syn1 antibody (Fig. 4C). In each sample the purity of the nuclear and cytoplasmic fractions was determined via their respective immunoreactivity towards Histone H3 and GAPDH (Fig. 4C). The levels of aSyn^Nuc^ were ~10 fold lower than those in the cytoplasmic fractions, independent of disease status (0.091±0.04 cf. 0.11±0.05 in controls cf. dementia with Lewy bodies respectively, p>0.05, data not shown). The levels of monomeric aSyn within the nuclear fraction, adjusted to nuclear loading control histone H3 were not significantly different between controls and dementia with Lewy bodies cases (Fig. 4D, p > 0.05). Longer exposures of Syn-1 blots, again revealed the occurrence of higher molecular weight aSyn species of ~28 kDa, 42-45 kDa and 54-56 kDa consistent with the presence of nuclear aSyn oligomers only in the nuclear extracts from cases of dementia with Lewy bodies (Fig. 4D, arrows). Importantly, the presence of identical levels of aSyn^Nuc^ monomer in nuclear fractions derived from control and cases of dementia with Lewy bodies, argues against the detection of aSyn merely as a consequence of the erroneous enrichment of aggregated aSyn due to the fractionation processes. Similar results were also obtained using MJRF1 pan-aSyn antibody (Supplementary Fig. 2). We also found that both monomeric and oligomeric aSyn^Nuc^ were phosphorylated at Ser-129 and the levels of pS129 reactive monomers were elevated in cases of dementia with Lewy bodies compared to controls (14.3±3.2-fold increase, p<0.01, Fig. 4D). Critically, no immunoreactivity towards Syn-1, MJFR1 or pS129 was detected in similarly processed tissue from aSyn knockout mice, confirming the specificity of our immunoreactivity (Fig. 4 and Supplementary Fig. 2). To further rule out the potential of cytoplasmic contamination of the nuclear fraction underlying the detection of aSyn^Nuc^, comparative analysis of GAPDH expression across cytoplasmic and nuclear fraction was conducted. Of the 13 cases investigated a quantifiable GAPDH signal within the nuclear fraction was detected in only 6 samples (~46%), at a level ~300 fold lower than that of cytoplasmic GAPDH (Supplementary Fig. 3). The magnitude of cytoplasmic contamination within the nuclear preparations was below that of the level of aSyn^Nuc^ detected in comparison to cytoplasmic aSyn and thus cannot account for the present of aSyn within the nuclear fraction.

### Label-free detection of aSyn within isolated nuclear preparations

Whilst the above data strongly supports the genuine nuclear presence of aSyn, this was entirely dependent on indirect antibody conjugation and, therefore, we could not rule out the occurrence of non-aSyn protein(s) in human tissue which share considerable cross-reactivity with commonly used aSyn antibodies. To address this limitation, both cytoplasmic and nuclear fractions were further investigated via mass spectrometry. Pooling of peptide sequence data across controls and cases of dementia with Lewy bodies enabled the generation of nuclear (~7500 identified proteins) and cytoplasmic (~5700 identified proteins) proteomic libraries. Enrichment analysis confirmed distinct proteomic profiles between the two fractions, cytoplasmic samples being clearly segregated from nuclear samples via principle component analysis (see Fig. 5A) and specific proteins reported as selectively enriched in either one or the other fraction (see Fig. 5B). Fractions were further validated for nuclear and cytoplasmic cellular compartment enrichments as per ShinyGo analysis (see Fig. 5C for top 10 hits and Supplementary Table 2, for top 30 hits). All three synucleins (alpha-, beta-, and gamma-synuclein) were detected within the nuclear fraction, consistent with data presented above and prior work of other groups^15,38^ (Table 3). With respect to aSyn, 7 peptides were detected, 6 of which were unique peptides only possible to have originated from aSyn (NP_000336.1) (combined, these peptides provided 71.4% aSyn sequence coverage within the nuclear fraction). The 6 unique peptides cover two large sequences of the aSyn protein: aa44-aa96, which includes the aggregation prone non-amyloid β component region, and aa103-aa140, encompassing the disorganised C-terminal (See Fig. 5D for peptide sequences and their alignment and Supplementary Fig. 4 for spectral traces). For comparison 15 peptides, 12 of which were unique and together provided 87.9% sequence coverage of the cytoplasmic fraction (Fig. 5D).

**Figure 5.**
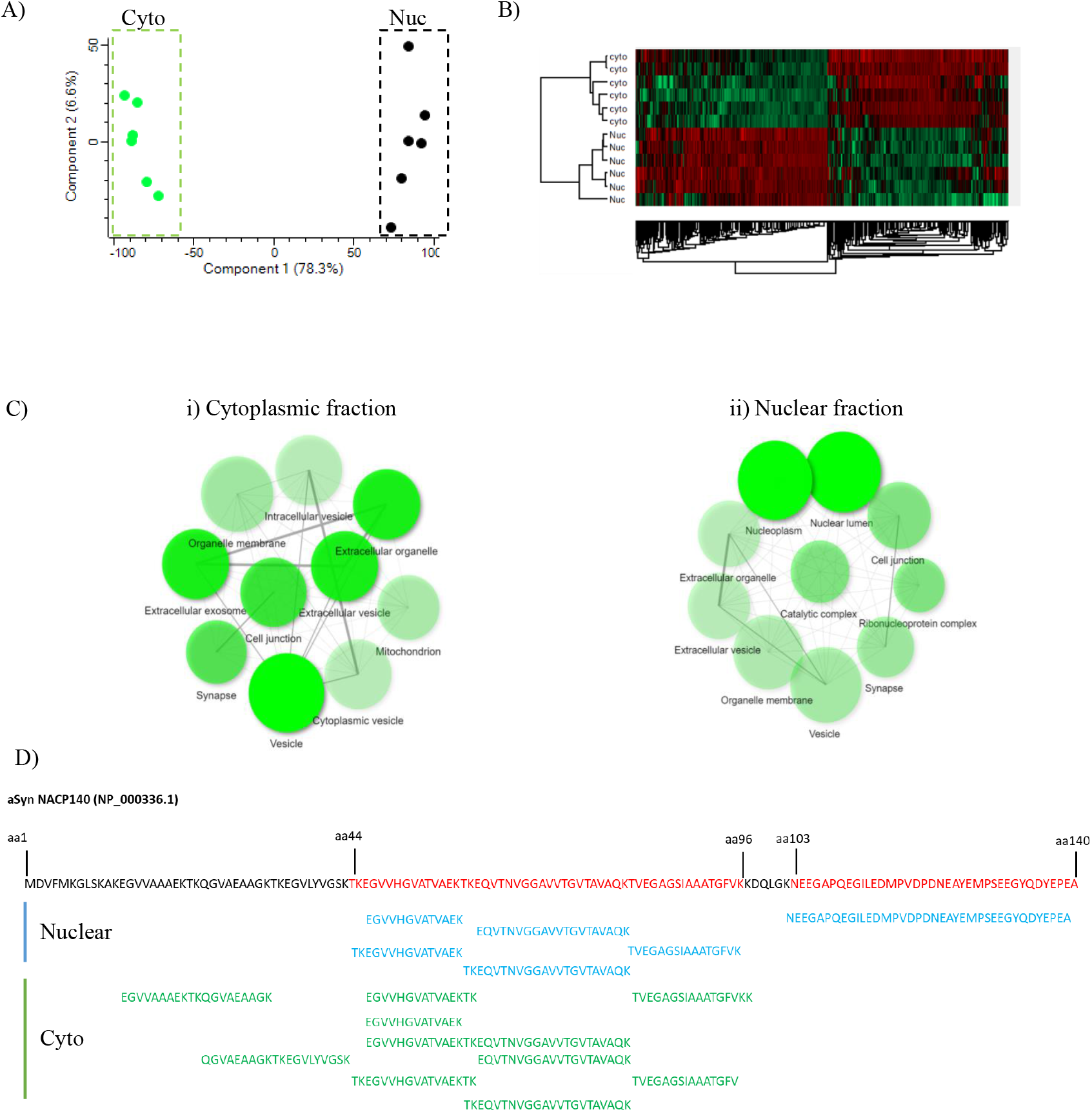
Mass spectrometry fractional analysis and aSyn peptide sequence alignment. **A**) Principal component analysis (PCA) of the relative abundance of proteins across pooled protein library, demonstrated clear distinction between cytoplasmic (Cyto) and Nuclear (Nuc) fractions. **B**). Such a segregation was equally apparent when individual protein comparison between the fractions of relative protein abundance was expressed as a heat map. **C**) Gene ontology cell component enrichment analysis of mass-spectrometry identified proteosome from nuclear (i) and cytoplasmic fraction (ii). Cell component networks are shown as produced from Shiny GO (V 0.66). For each node, darker shading represents significantly greater gene enrichment, node size indicates the number of genes aligning with the specific cellular component node and the thickness of connecting lines demonstrates the magnitude of overlapping genes between nodes. **D**) Peptide mapping to the aSyn amino acid (aa) sequence, unique peptides recovered from mass spectrometry of nuclear fraction are shown in blue and those recovered from the cytoplasmic fraction shown in green. Full length aSyn sequence (NP_000336.1) is shown above with the sequence covered by peptides present in nuclear fraction shown in red.

**Table 3.** Mass Spectrometry detection of synuclein-family members. Number of peptides, sequence coverage are provided. For number of peptides and sequence coverage, values are given as per total, unique peptide and unique peptide following razor peptide adjustment.

| Protein | Peptides |  |  | Sequence coverage [%] |  |  |
| --- | --- | --- | --- | --- | --- | --- |
|  | Total | Unique | Unique + Razor | Total | Unique | Unique + Razor |
| Nuclear |  |  |  |  |  |  |
| Alpha-synuclein | 7 | 6 | 7 | 71.4 | 65 | 71.4 |
| Beta-synuclein | 4 | 3 | 3 | 36.6 | 29.9 | 29.9 |
| Gamma-synuclein | 6 | 6 | 6 | 44.9 | 44.9 | 44.9 |
| Cytoplasmic |  |  |  |  |  |  |
| Alpha-synuclein | 15 | 12 | 12 | 87.9 | 87.9 | 87.9 |
| Beta-synuclein | 17 | 14 | 17 | 64.2 | 64.2 | 64.2 |
| Gamma-synuclein | 19 | 19 | 19 | 95.3 | 95.3 | 95.3 |

## Discussion

In this study, we have demonstrated the occurrence of aSyn in the nucleus of cells in human brain tissue based not only by indirect in-situ antibody labelling and biochemical analysis, but also via label free mass-spectrometry. These data strongly support the physiological occurrence of aSyn^Nuc^ in human brain tissue as nuclear detection was apparent in all cases investigated, independent of disease status. Furthermore, our study highlights a novel nuclear-centric aSyn pathology which occurs alongside the cytoplasmic formation of Lewy bodies.

### Presence of aSyn in the nucleus

Although we are the first to systematically characterise aSyn^Nuc^ in human tissue, aSyn^Nuc^ has been reported at endogenous levels in brain tissue from a variety of animal species, including mice and primates^12,17,18,22,39 40^ and, originally, in electric ray^10^ as well being detected in various neuronal culture preparations.^17,19,41^ Likewise, aSyn^Nuc^ has been reported in transgenic aSyn overexpression animals^42^ as well as in overexpression cell models.^14,15,43–45^ In these model systems, increased expression of aSyn presumably enhanced the levels of aSyn^Nuc^ enabling easier detection, some of which have avoid the concerns of antibody cross reactivity by the implementation of either fluorescent or reporter fusion proteins.^44,45^ Combined the collective evidence supports a conserved physiological occurrence of aSyn^Nuc^ in the brain. Nevertheless, given that several commercial aSyn antibodies demonstrate non-aSyn specific cross-reactivity,^11^ and that many neuropathological reports have failed to observe aSyn^Nuc^ staining in human tissue,^1,25,27,46^ this issue has remained controversial. Here, we demonstrate that the specific antigen retrieval methods employed upon fixed tissue prior to immunohistochemical protocols is a key factor in the detection of aSyn^Nuc^. As common and recommended antigen retrieval methods for the quantification of LB pathology, namely formic acid and proteinase K,^47^ were suboptimal for aSyn^Nuc^ detection, it is likely that the occurrence of aSyn^Nuc^ in the human brain has been largely overlooked and underreported as a consequence of the poor nuclear antigen access gained when employing such antigen retrieval methods. Thus, the use of antigen retrieval methods optimised for nuclear antigens, such as pressurised heating in EDTA can be considered critical for a determination of nuclear pathology, particularly when using human brain tissue where the fixation duration can be considerably longer than in cell and animal work. Likewise, considering the low levels of endogenous aSyn^Nuc^ in human brain tissue, technical limitations may have previously precluded immunoblot quantification of aSyn^Nuc^. Loss of low molecular weight proteins, including aSyn, from transfer membranes has previously been reported and can lead to the concentration of target proteins to fall below detectable levels without post-transfer membrane fixation.^37^ In the present study, we evaluated a paraformaldehyde post-transfer fixation method, previously evaluated for increased aSyn retention^37^ as well as a fixation method of boiling the membrane in phosphate buffered saline, which is more commonly used for the detection of monomeric β-amyloid.^48^ Although the specifics of this process are still under investigation,^49^ it can be assumed that optimal conditions for the membrane retention of a particular protein of interest varies depending on chemical composition. For this reason, it is interesting to note that whilst heating in phosphate buffered saline was more favourable for optimising the signal of aSyn^Nuc^, paraformaldehyde was more favourable for monomeric cytoplasmic aSyn, suggesting a difference in chemical composition between cytoplasmic and nuclear pools of aSyn.

Such differences are likely the result of chemical post-translational modifications as opposed to truncations, as despite some evidence of Syn1 antibody-reactive truncation products via immunoblot, the majority of reactivity was in line with the molecular weight of full length aSyn. Moreover, the positive aSyn^Nuc^ histochemical staining following aa2-12 (Syn-303), aa91-99 (Syn-1) and aa129 (pS129) directed antibodies in addition to the peptide sequences recovered from mass-spectrometry covering aa44-96 and aa103-140, aSyn^Nuc^, further supports the nuclear import of the full-length protein. This is consistent with the increased nuclear localisation of full length aSyn compared to various deletion mutants, despite the nuclear exclusion properties of the central region of the protein, including of the non-amyloid β component region.^15^ The lack of unique N-terminal generated peptides associated with aSyn presumably is as a consequence of the highly conserved amino acid sequence of the N-terminus between α, β and γ-Syn.^50^

### Physiological localization of aSyn^Nuc^

The occurrence of aSyn^Nuc^ in human brain tissue appears widespread, as aSyn positive nuclei were frequently observed and aSyn^Nuc^ was present in both immunohistochemistry and immunoblot experiments, in all cases examined, independent of disease status. Moreover, the detection of pS129 aSyn was not limited to NeuN positive neurons but also present in the nucleus within NeuN negative cells likely of glial lineage. Although largely considered a neuronal specific protein, *in-vitro* cell culture studies^51–53^ and human brain immunohistochemistry^54^ supports the low-level expression of aSyn within astrocytes and oligodendrocytes in physiological conditions. Such ubiquity of aSyn^Nuc^ implies its involvement in physiological processes. Accordingly, recent studies have associated aSyn^Nuc^ with genomic integrity, either directly by participating in DNA double strand break repair,^17^ or by the enhanced transcription of numerous genes including those associated DNA damage and repair via the modulation of retinoic acid mediated gene expression.^19^ Critically, however, when overexpressed, aSyn^Nuc^ widely downregulates gene transcription either as a consequence of inhibited histone acetylation,^13,22^ or via direct interactions with gene promotor regions.^24^ Among the transcripts affected, those associated with cell cycle, DNA repair^13,14^ and mitochondrial regulation^24^ clearly highlight the potential for dysfunctional aSyn^Nuc^ to engage in several well-established neurodegenerative pathways known to contribute the cellular pathology associated with dementia with Lewy bodies.

### Disease-dependent modification of aSyn^Nuc^

We observed aSyn^Nuc^ pathology in cases of dementia with Lewy bodies, comprising hyperphosphorylated serine 129 aSyn and higher molecular weight SDS-stable oligomeric aSyn species. Both phosphorylation^55,56^ and oliogmerisation^57^ of aSyn are widely associated with cytoplasmic Lewy body pathology and, therefore, the nuclear pathology is consistent with speculated pathological mechanisms.^58^ However, identical modified molecular weight banding of aSyn were not detected in the cytoplasmic fraction of matched cases of dementia with Lewy bodies, suggesting the changes in aSyn^Nuc^ composition are unlikely to solely reflect changes in whole cell aSyn composition and likely represents novel conformers of aSyn that are distinct to the nucleus.

In both control and cases of dementia with Lewy bodies, aSyn^Nuc^ was reactive to phosphorylation-specific antibodies, albeit in dementia with Lewy bodies this was ~14-fold higher. Accordingly, it can be assumed that phosphorylation of aSyn^Nuc^, at least at low levels, is not overtly pathological. Indeed, cellular studies imply pS129aSyn^Nuc^ can be regulated endogenously in neurons by Polo-like 2 kinase activity^59^ and this promotes the nuclear import of aSyn.^14,60^ In the nucleus, phosphorylation may further regulate the activity of aSyn^Nuc^, by facilitating the recruitment of aSyn^Nuc^ to sites of DNA damage, modulating its transcriptional effects and promoting protection against cellular stress,^14,17^ whilst also preventing the formation of intranuclear oligomers.^20^

Nevertheless, despite these potentially beneficial roles for phosphorylated aSyn^Nuc^, some studies also point to the increased nuclear retention of pS129 aSyn,^44^ conducive to the protein’s subcellular accumulation beyond a physiological range, which may in turn be detrimental. This is particularly evident from *in-vivo* studies, where enhanced aSyn^Nuc^ via a nuclear localisation signal results in the loss of dopaminergic neurons and motor impairments in drosophila^22^ and mice,^61^ and by the cellular toxicity associated with the nuclear translocation of aSyn in response to oxidative stressors.^18,24,62,63^ Although unresolved contradictions regarding the toxicity of aSyn with a nuclear localisation signal^14^ and oxidative stress induced aSyn^Nuc^ accumulation^64^ exist, and thus may reflect differences in expression levels, cell type specific vulnerability, or the differential generation of aSyn^Nuc^ species.

As a consequence of increased compartmental concentration and, potentially, of altered post-translational modification, the risk of self-aggregation for aSyn^Nuc^ in disease conditions is likely altered/increased.^9^ The observed pS129 positive oligomeric aSyn species would support this, and are consistent with our previous observations of intranuclear reactivity towards oligomer/aggregation specific aSyn antibodies in cases of dementia with Lewy bodies and the formation of aSyn^Nuc^ oligomers in cellular overexpression systems.^14^ Certainly, the nuclear environment would appear prone to aggregation, as both DNA^65–67^ and histones^18^ appear to accelerate the aggregation process of aSyn, and other amyloidogenic proteins, into β-sheet fibrils. Although we did not detect any overt signs of mature fibril-like inclusions in isolated nuclei, neuronal and oligodendrocytic intranuclear inclusions of β-sheet rich aSyn filaments are a pathological feature of multiple system atrophy, alongside aSyn cytoplasmic aggregates.^16^ As part of the disease process of multiple system atrophy, elevated pS129 aSyn^Nuc^ is considered as an early pathological event, present in preclinical cases^68^ and patient derived neural progenitor cells.^69^ This initial increase in intranuclear diffuse or punctate staining, similar to what was observed in the present study, is assumed to precede the formation of thioflavin positive aggregates.^68^ Given the emerging evidence suggesting the innate differences in disease specific aSyn strains,^70–72^ it is possible that, across synucleiopathies, the homeostasis of aSyn^Nuc^ may be commonly affected, yet the propensity for the formation of intranuclear aggregation may differ between diseases. In any case, the occurrence of neuronal nuclear inclusions in the brain stem nuclei of multiple system atrophy cases correlates with surviving neuron numbers, suggestive of a protective role for mature inclusions,^73^ which is in line with cellular toxicity being largely independent of overt aSyn aggregates in cells models^62^ and that instead, at least within the nucleus, toxicity may be driven by intermediate aSyn oligomers.^74^

## Conclusion

The present study has addressed an outstanding question in the field by confirming the nuclear occurrence of aSyn using an array of histological and molecular methods in human *post-mortem* tissue. Furthermore, we have identified that aSyn^Nuc^ is significantly increased in cases of dementia with Lewy bodies and manifests increased levels of putatively pathogenic post-translation modifications, such as pS129. Intriguingly, aSyn^Nuc^ is enriched for oligomeric species distinct from those observed in the cytoplasm and, given the potential role of oligomers in driving disease-associated toxicity, such species may result in toxicity that drives cellular impairment and neurodegeneration associated with dementia with Lewy bodies.

## Abbreviations

aSyn: Alpha-synuclein
βSyn: Beta-synuclein
γSyn: Gamma-synuclein
aSyn^Nuc^: Nuclear alpha-synuclein
CERAD: Consortium to establish a registry for Alzheimer’s disease
DAPI: 4’,6-diamidino-2-phenylindole
FDR: False discovery rate
GAPDH: Glyceraldehyde 3-phosphate dehydrogenase
LCMS: Liquid chromatography mass spectrometry
NIA-AA: National institute of ageing-Alzheimer’s Association
SDS: Sodium dodecyl sulfate

## Acknowledgements

Tissue for this study was provided by the Newcastle Brain Tissue Resource which is funded in part by a grant from the UK Medical Research Council (G0400074), by NIHR Newcastle Biomedical Research Centre awarded to the Newcastle upon Tyne NHS Foundation Trust and Newcastle University, and as part of the Brains for Dementia Research Programme jointly funded by Alzheimer’s Research UK and Alzheimer’s Society.

## Supplementary Table legends

**Supplemental Table 1. Individual Human cases.** Diagnosis (Diag), age (in years), sex, post mortem interval (PMI, in hrs) and neuropathological assessment scores for neurofibrillary tangle (NFT) Braak stage, Thal phase, Consortium to Establish a Registry for Alzheimer’s Disease (CERAD), the National Institute of Ageing – Alzheimer’s Association (NIA-AA) criteria, Lewy body (LB) Braak stage and McKeith criteria are provided. For McKeith criteria, absence of Lewy pathology (No LB), Limbic predominate and Neocortical (neoctrx) predominate are indicated. Additionally, the use of each case in western blots (WB), multichannel fluorescence histochemistry (His) and/or mass spectrometry (MS) is also listed. N.A= not available.

**Supplemental Table 2. Fraction enrichment.** Top 30 cellular component categories identified from proteomic analysis of nuclear and cytoplasmic tissue fractions as per ShinyGo database. Assigned category name, number of proteins associated genes detected out of total list are provided as well as false discovery rate (FDR).

## Supplementary Figure legends

**Supplemental Figure 1. Nuclear antigen accessibility following antigen retrieval methods**. Representative images captured via a 40x objective lens from control cases stained for Histone H3 with DAPI nuclear stained. Comparison of staining with citrate, formic acid and EDTA + formic acid clearly demonstrates optimum nuclear labelling following a combined treatment of EDTA and formic acid. Scale bar = 10 μm.

**Supplemental Figure 2. Detection of nuclear aSyn via pan-aSyn antibody MJRF1**. Example western blots of pan-aSyn MJFR1 immunoreactivity of cytoplasmic (c) and nuclear (N) fractionates from control (con) and cases of dementia with Lewy bodies (DLB) (1.8μg/lane). Monomeric aSyn is shown under optimised exposure conditions, with large panel depicting monomeric and oligomeric aSyn species captured following overexposure of the blot. Antibody specificity was confirmed by means of similar probing of tissue fractionates generated from aSyn knockout mice (aSyn). Cytoplasmic and nuclear loading controls GAPDH and Histone H3 are also shown. Note faint appearance of high molecular weight aSyn species in the nuclear fraction only in DLB cases (arrows) only in the nuclear fraction.

**Supplemental Figure 3. Quantification of cytoplasmic proteins within nuclear fractions**. Example western blots of GAPDH immunoreactivity of cytoplasmic (C) and nuclear (N) fractionates (**A**) without (i) and with (ii) enhanced contrast to allow for visualisation of GAPDH within the nuclear fraction. Comparative analysis between GAPDH immunoreactivity from cytoplasmic and those sample in which GAPDH was above detection threshold (~46%) indicated ~ a 300 fold dilution of cytoplasmic components in the nuclear fraction (**B**).

**Supplemental Figure 4. Mass spectrometry detection of aSyn in the nuclear fraction**. Annotated spectra for each of the recovered peptides, b and y ions are highlighted accordingly.

## References

1. Beach TG, Adler CH, Lue L, et al. Unified staging system for Lewy body disorders: correlation with nigrostriatal degeneration, cognitive impairment and motor dysfunction. Acta neuropathologica. Jun 2009;117(6):613–34. doi:10.1007/s00401-009-0538-8

2. Milber JM, Noorigian JV, Morley JF, et al. Lewy pathology is not the first sign of degeneration in vulnerable neurons in Parkinson disease. Neurology. Dec 11 2012;79(24):2307–14. doi:10.1212/WNL.0b013e318278fe32

3. Dijkstra AA, Voorn P, Berendse HW, et al. Stage-dependent nigral neuronal loss in incidental Lewy body and Parkinson’s disease. Movement disorders: official journal of the Movement Disorder Society. 2014;29(10):1244–1251. doi:10.1002/mds.25952

4. Katsuse O, Iseki E, Marui W, Kosaka K. Developmental stages of cortical Lewy bodies and their relation to axonal transport blockage in brains of patients with dementia with Lewy bodies. Journal of the neurological sciences. Jul 15 2003;211(1-2):29–35.

5. Serrano-Pozo A, Qian J, Muzikansky A, et al. Thal Amyloid Stages Do Not Significantly Impact the Correlation Between Neuropathological Change and Cognition in the Alzheimer Disease Continuum. Journal of neuropathology and experimental neurology. 2016;75(6):516–526. doi:10.1093/jnen/nlw026

6. Rudinskiy N, Hawkes JM, Wegmann S, et al. Tau pathology does not affect experience-driven single-neuron and network-wide Arc/Arg3.1 responses. Acta neuropathologica communications. 2014;2:63–63. doi:10.1186/2051-5960-2-63

7. Polymeropoulos MH, Lavedan C, Leroy E, et al. Mutation in the alpha-synuclein gene identified in families with Parkinson’s disease. Science. Jun 27 1997;276(5321):2045–7.

8. Deng H, Yuan L. Genetic variants and animal models in SNCA and Parkinson disease. Ageing Res Rev. May 2014;15:161–76. doi:10.1016/j.arr.2014.04.002

9. Marvian AT, Koss DJ, Aliakbari F, Morshedi D, Outeiro TF. In vitro models of synucleinopathies: informing on molecular mechanisms and protective strategies. Journal of neurochemistry. Sep 2019;150(5):535–565. doi:10.1111/jnc.14707

10. Maroteaux L, Campanelli JT, Scheller RH. Synuclein: a neuron-specific protein localized to the nucleus and presynaptic nerve terminal. The Journal of neuroscience: the official journal of the Society for Neuroscience. 1988;8(8):2804–2815. doi:10.1523/JNEUROSCI.08-08-02804.1988

11. Huang Z, Xu Z, Wu Y, Zhou Y. Determining nuclear localization of alpha-synuclein in mouse brains. Neuroscience. 2011;199:318–332. doi:10.1016/j.neuroscience.2011.10.016

12. Yu S, Li X, Liu G, et al. Extensive nuclear localization of alpha-synuclein in normal rat brain neurons revealed by a novel monoclonal antibody. Neuroscience. 2007;145(2):539–555. doi:10.1016/j.neuroscience.2006.12.028

13. Paiva I, Pinho R, Pavlou MA, et al. Sodium butyrate rescues dopaminergic cells from alpha-synuclein-induced transcriptional deregulation and DNA damage. Human molecular genetics. 2017;26(12):2231–2246. doi:10.1093/hmg/ddx114

14. Pinho R, Paiva I, Jercic KG, et al. Nuclear localization and phosphorylation modulate pathological effects of alpha-synuclein. Human molecular genetics. 2019;28(1):31–50. doi:10.1093/hmg/ddy326

15. Specht CG, Tigaret CM, Rast GF, Thalhammer A, Rudhard Y, Schoepfer R. Subcellular localisation of recombinant alpha- and gamma-synuclein. Molecular and cellular neurosciences. Feb 2005;28(2):326–34. doi:10.1016/j.mcn.2004.09.017

16. Lin WL, DeLucia MW, Dickson DW. Alpha-synuclein immunoreactivity in neuronal nuclear inclusions and neurites in multiple system atrophy. Neuroscience letters. Jan 9 2004;354(2):99–102. doi:10.1016/j.neulet.2003.09.075

17. Schaser AJ, Osterberg VR, Dent SE, et al. Alpha-synuclein is a DNA binding protein that modulates DNA repair with implications for Lewy body disorders. Scientific reports. 2019;9(1):10919–10919. doi:10.1038/s41598-019-47227-z

18. Goers J, Manning-Bog AB, McCormack AL, et al. Nuclear localization of alpha-synuclein and its interaction with histones. Biochemistry. 2003;42(28):8465–8471. doi:10.1021/bi0341152

19. Davidi D, Schechter M, Elhadi SA, Matatov A, Nathanson L, Sharon R. α-Synuclein Translocates to the Nucleus to Activate Retinoic-Acid-Dependent Gene Transcription. iScience. Mar 27 2020;23(3):100910. doi:10.1016/j.isci.2020.100910

20. Lazaro DF, Rodrigues EF, Langohr R, et al. Systematic comparison of the effects of alpha-synuclein mutations on its oligomerization and aggregation. PLoS genetics. Nov 2014;10(11):e1004741. doi:10.1371/journal.pgen.1004741

21. Fares MB, Ait-Bouziad N, Dikiy I, et al. The novel Parkinson’s disease linked mutation G51D attenuates in vitro aggregation and membrane binding of α-synuclein, and enhances its secretion and nuclear localization in cells. Hum Mol Genet. Sep 1 2014;23(17):4491–509. doi:10.1093/hmg/ddu165

22. Kontopoulos E, Parvin JD, Feany MB. Alpha-synuclein acts in the nucleus to inhibit histone acetylation and promote neurotoxicity. Human molecular genetics. 2006;15(20):3012–3023. doi:10.1093/hmg/ddl243

23. Chen V, Moncalvo M, Tringali D, et al. The mechanistic role of alpha-synuclein in the nucleus: impaired nuclear function caused by familial Parkinson’s disease SNCA mutations. Hum Mol Genet. Nov 4 2020;29(18):3107–3121. doi:10.1093/hmg/ddaa183

24. Siddiqui A, Chinta SJ, Mallajosyula JK, et al. Selective binding of nuclear alpha-synuclein to the PGC1alpha promoter under conditions of oxidative stress may contribute to losses in mitochondrial function: implications for Parkinson’s disease. Free Radic Biol Med. Aug 15 2012;53(4):993–1003. doi:10.1016/j.freeradbiomed.2012.05.024

25. Braak H, Del Tredici K, Rub U, de Vos RA, Jansen Steur EN, Braak E. Staging of brain pathology related to sporadic Parkinson’s disease. Neurobiology of aging. Mar-Apr 2003;24(2):197–211.

26. McKeith IG, Boeve BF, Dickson DW, et al. Diagnosis and management of dementia with Lewy bodies: Fourth consensus report of the DLB Consortium. Review. Neurology. Jul 4 2017;89(1):88–100. doi:10.1212/WNL.0000000000004058

27. McKeith IG, Dickson DW, Lowe J, et al. Diagnosis and management of dementia with Lewy bodies: third report of the DLB Consortium. Neurology. Dec 27 2005;65(12):1863–72. doi:10.1212/01.wnl.0000187889.17253.b1

28. Montine TJ, Phelps CH, Beach TG, et al. National Institute on Aging-Alzheimer’s Association guidelines for the neuropathologic assessment of Alzheimer’s disease: a practical approach. Acta neuropathologica. Jan 2012;123(1):1–11. doi:10.1007/s00401-011-0910-3

29. Braak H, Alafuzoff I, Arzberger T, Kretzschmar H, Del Tredici K. Staging of Alzheimer disease-associated neurofibrillary pathology using paraffin sections and immunocytochemistry. Acta neuropathologica. Oct 2006;112(4):389–404. doi:10.1007/s00401-006-0127-z

30. Thal DR, Rüb U, Orantes M, Braak H. Phases of A beta-deposition in the human brain and its relevance for the development of AD. Neurology. Jun 25 2002;58(12):1791–800. doi:10.1212/wnl.58.12.1791

31. Mirra SS, Heyman A, McKeel D, et al. The Consortium to Establish a Registry for Alzheimer’s Disease (CERAD). Part II. Standardization of the neuropathologic assessment of Alzheimer’s disease. Neurology. Apr 1991;41(4):479–86. doi:10.1212/wnl.41.4.479

32. Robertson DC, Schmidt O, Ninkina N, Jones PA, Sharkey J, Buchman VL. Developmental loss and resistance to MPTP toxicity of dopaminergic neurones in substantia nigra pars compacta of gamma-synuclein, alpha-synuclein and double alpha/gamma-synuclein null mutant mice. Journal of neurochemistry. Jun 2004;89(5):1126–36. doi:10.1111/j.1471-4159.2004.02378.x

33. Majbour NK, Vaikath NN, van Dijk KD, et al. Oligomeric and phosphorylated alpha-synuclein as potential CSF biomarkers for Parkinson’s disease. Molecular neurodegeneration. Jan 19 2016;11:7. doi:10.1186/s13024-016-0072-9

34. Matevossian A, Akbarian S. Neuronal nuclei isolation from human postmortem brain tissue. J Vis Exp. 2008;(20):914. doi:10.3791/914

35. Ge SX, Jung D, Yao R. ShinyGO: a graphical gene-set enrichment tool for animals and plants. Bioinformatics. Apr 15 2020;36(8):2628–2629. doi:10.1093/bioinformatics/btz931

36. Tyanova S, Temu T, Sinitcyn P, et al. The Perseus computational platform for comprehensive analysis of (prote)omics data. Nature methods. 2016/09/01 2016;13(9):731–740. doi:10.1038/nmeth.3901

37. Lee BR, Kamitani T. Improved immunodetection of endogenous α-synuclein. PloS one. 2011;6(8):e23939. doi:10.1371/journal.pone.0023939

38. Tanji K, Mori F, Nakajo S, et al. Expression of beta-synuclein in normal human astrocytes. Neuroreport. Sep 17 2001;12(13):2845–8. doi:10.1097/00001756-200109170-00018

39. Mori F, Tanji K, Yoshimoto M, Takahashi H, Wakabayashi K. Immunohistochemical comparison of alpha- and beta-synuclein in adult rat central nervous system. Brain research. Jun 21 2002;941(1-2):118–26. doi:10.1016/s0006-8993(02)02643-4

40. McCormack AL, Mak SK, Di Monte DA. Increased α-synuclein phosphorylation and nitration in the aging primate substantia nigra. Cell Death Dis. May 31 2012;3(5):e315. doi:10.1038/cddis.2012.50

41. Ryu S, Baek I, Liew H. Sumoylated α-synuclein translocates into the nucleus by karyopherin α6. Molecular & Cellular Toxicology. 2019/01/01 2019;15(1):103–109. doi:10.1007/s13273-019-0012-1

42. Masliah E, Rockenstein E, Veinbergs I, et al. Dopaminergic loss and inclusion body formation in alpha-synuclein mice: implications for neurodegenerative disorders. Science. Feb 18 2000;287(5456):1265–9. doi:10.1126/science.287.5456.1265

43. Yuan Y, Jin J, Yang B, et al. Overexpressed alpha-synuclein regulated the nuclear factor-kappaB signal pathway. Cell Mol Neurobiol. Jan 2008;28(1):21–33. doi:10.1007/s10571-007-9185-6

44. Gonçalves S, Outeiro TF. Assessing the subcellular dynamics of alpha-synuclein using photoactivation microscopy. Mol Neurobiol. Jun 2013;47(3):1081–92. doi:10.1007/s12035-013-8406-x

45. McLean PJ, Ribich S, Hyman BT. Subcellular localization of alpha-synuclein in primary neuronal cultures: effect of missense mutations. J Neural Transm Suppl. 2000;(58):53–63. doi:10.1007/978-3-7091-6284-2_5

46. Leverenz JB, Hamilton R, Tsuang DW, et al. Empiric refinement of the pathologic assessment of Lewy-related pathology in the dementia patient. Brain pathology (Zurich, Switzerland). Apr 2008;18(2):220–4. doi:10.1111/j.1750-3639.2007.00117.x

47. Beach TG, White CL, Hamilton RL, et al. Evaluation of alpha-synuclein immunohistochemical methods used by invited experts. Acta neuropathologica. Sep 2008;116(3):277–88. doi:10.1007/s00401-008-0409-8

48. Koss DJ, Jones G, Cranston A, Gardner H, Kanaan NM, Platt B. Soluble pre-fibrillar tau and ß-amyloid species emerge in early human Alzheimer’s disease and track disease progression and cognitive decline. Acta neuropathologica. Dec 2016;132(6):875–895. doi:10.1007/s00401-016-1632-3

49. Xu J, Sun H, Huang G, et al. A fixation method for the optimisation of western blotting. Scientific reports. Apr 30 2019;9(1):6649. doi:10.1038/s41598-019-43039-3

50. George JM. The synucleins. Genome Biol. 2002;3(1):Reviews3002. doi:10.1186/gb-2001-3-1-reviews3002

51. Tanji K, Imaizumi T, Yoshida H, et al. Expression of alpha-synuclein in a human glioma cell line and its up-regulation by interleukin-1beta. Neuroreport. Jul 3 2001;12(9):1909–12. doi:10.1097/00001756-200107030-00028

52. Gu XL, Long CX, Sun L, Xie C, Lin X, Cai H. Astrocytic expression of Parkinson’s disease-related A53T alpha-synuclein causes neurodegeneration in mice. Molecular brain. Apr 21 2010;3:12. doi:10.1186/1756-6606-3-12

53. Richter-Landsberg C, Gorath M, Trojanowski JQ, Lee VM. alpha-synuclein is developmentally expressed in cultured rat brain oligodendrocytes. J Neurosci Res. Oct 1 2000;62(1):9–14. doi:10.1002/1097-4547(20001001)62:1<9::Aid-jnr2>3.0.Co;2-u

54. Mori F, Tanji K, Yoshimoto M, Takahashi H, Wakabayashi K. Demonstration of alpha-synuclein immunoreactivity in neuronal and glial cytoplasm in normal human brain tissue using proteinase K and formic acid pretreatment. Experimental neurology. Jul 2002;176(1):98–104. doi:10.1006/exnr.2002.7929

55. Fujiwara H, Hasegawa M, Dohmae N, et al. alpha-Synuclein is phosphorylated in synucleinopathy lesions. Nature cell biology. Feb 2002;4(2):160–4. doi:10.1038/ncb748

56. Anderson JP, Walker DE, Goldstein JM, et al. Phosphorylation of Ser-129 is the dominant pathological modification of alpha-synuclein in familial and sporadic Lewy body disease. The Journal of biological chemistry. 2006;281(40):29739–29752. doi:10.1074/jbc.M600933200

57. Grassi D, Howard S, Zhou M, et al. Identification of a highly neurotoxic alpha-synuclein species inducing mitochondrial damage and mitophagy in Parkinson’s disease. Proceedings of the National Academy of Sciences of the United States of America. Mar 13 2018;115(11):E2634–e2643. doi:10.1073/pnas.1713849115

58. Outeiro TF, Koss DJ, Erskine D, et al. Dementia with Lewy bodies: an update and outlook. Molecular neurodegeneration. Jan 21 2019;14(1):5. doi:10.1186/s13024-019-0306-8

59. Elfarrash S, Jensen NM, Ferreira N, et al. Polo-like kinase 2 inhibition reduces serine-129 phosphorylation of physiological nuclear alpha-synuclein but not of the aggregated alpha-synuclein. bioRxiv. 2021:2021.05.21.445104. doi:10.1101/2021.05.21.445104

60. Schell H, Hasegawa T, Neumann M, Kahle PJ. Nuclear and neuritic distribution of serine-129 phosphorylated alpha-synuclein in transgenic mice. Neuroscience. Jun 2 2009;160(4):796–804. doi:10.1016/j.neuroscience.2009.03.002

61. Ma KL, Song LK, Yuan YH, et al. The nuclear accumulation of alpha-synuclein is mediated by importin alpha and promotes neurotoxicity by accelerating the cell cycle. Neuropharmacology. Jul 2014;82:132–42. doi:10.1016/j.neuropharm.2013.07.035

62. Villar-Piqué A, Rossetti G, Ventura S, Carloni P, Fernández CO, Outeiro TF. Copper(II) and the pathological H50Q α-synuclein mutant: Environment meets genetics. Commun Integr Biol. 2017;10(1):e1270484. doi:10.1080/19420889.2016.1270484

63. Zhou M, Xu S, Mi J, Uéda K, Chan P. Nuclear translocation of alpha-synuclein increases susceptibility of MES23.5 cells to oxidative stress. Brain research. Mar 15 2013; 1500:19–27. doi:10.1016/j.brainres.2013.01.024

64. Liu X, Lee YJ, Liou LC, et al. Alpha-synuclein functions in the nucleus to protect against hydroxyurea-induced replication stress in yeast. Hum Mol Genet. Sep 1 2011;20(17):3401–14. doi:10.1093/hmg/ddr246

65. Cherny D, Hoyer W, Subramaniam V, Jovin TM. Double-stranded DNA stimulates the fibrillation of alpha-synuclein in vitro and is associated with the mature fibrils: an electron microscopy study. Journal of molecular biology. Dec 3 2004;344(4):929–38. doi:10.1016/j.jmb.2004.09.096

66. Hegde ML, Rao KS. DNA induces folding in alpha-synuclein: understanding the mechanism using chaperone property of osmolytes. Arch Biochem Biophys. Aug 1 2007;464(1):57–69. doi:10.1016/j.abb.2007.03.042

67. Cordeiro Y, Macedo B, Silva JL, Gomes MPB. Pathological implications of nucleic acid interactions with proteins associated with neurodegenerative diseases. Biophys Rev. Mar 2014;6(1):97–110. doi:10.1007/s12551-013-0132-0

68. Yoshida M. Multiple system atrophy: alpha-synuclein and neuronal degeneration. Neuropathology: official journal of the Japanese Society of Neuropathology. Oct 2007;27(5):484–93. doi:10.1111/j.1440-1789.2007.00841.x

69. Herrera-Vaquero M, Heras-Garvin A, Krismer F, et al. Signs of early cellular dysfunction in multiple system atrophy. Neuropathol Appl Neurobiol. Feb 2021;47(2):268–282. doi:10.1111/nan.12661

70. Peng C, Gathagan RJ, Covell DJ, et al. Cellular milieu imparts distinct pathological alpha-synuclein strains in alpha-synucleinopathies. Nature. May 2018;557(7706):558–563. doi:10.1038/s41586-018-0104-4

71. Candelise N, Schmitz M, Llorens F, et al. Seeding variability of different alpha synuclein strains in synucleinopathies. Ann Neurol. May 2019;85(5):691–703. doi:10.1002/ana.25446

72. Van der Perren A, Gelders G, Fenyi A, et al. The structural differences between patient-derived α-synuclein strains dictate characteristics of Parkinson’s disease, multiple system atrophy and dementia with Lewy bodies. Acta neuropathologica. Jun 2020;139(6):977–1000. doi:10.1007/s00401-020-02157-3

73. Nishie M, Mori F, Yoshimoto M, Takahashi H, Wakabayashi K. A quantitative investigation of neuronal cytoplasmic and intranuclear inclusions in the pontine and inferior olivary nuclei in multiple system atrophy. Neuropathol Appl Neurobiol. Oct 2004;30(5):546–54. doi:10.1111/j.1365-2990.2004.00564.x

74. Vasquez V, Mitra J, Hegde PM, et al. Chromatin-Bound Oxidized α-Synuclein Causes Strand Breaks in Neuronal Genomes in in vitro Models of Parkinson’s Disease. Journal of Alzheimer’s disease:JAD. 2017;60(s1):S133–S150. doi:10.3233/JAD-170342

