## Supplemental table 1 for "Nuclear alpha-synuclein is present in the human brain and is modified in dementia with Lewy bodies"

| case nr | Diag | age | sex | PMI | NFT Braak | Thal | CERAD | NIA-AA | LB Braak | McKeith | Use |
| --- | --- | --- | --- | --- | --- | --- | --- | --- | --- | --- | --- |
| 1 | Con | 73 | M | 25 | 0 | 0 | neg | not | 0 | no LB | WB |
| 2 | Con | 92 | M | 50 | III | 1 | neg | low | 0 | no LB | WB |
| 3 | Con | 81 | F | 75 | II | 4 | neg | low | 0 | no LB | WB |
| 4 | Con | 84 | M | 45 | II | 3 | neg | low | 0 | no LB | WB |
| 5 | Con | 81 | F | 19 | I | 3 | neg | low | 0 | no LB | WB |
| 6 | Con | 92 | M | 9 | II | 3 | neg | low | 0 | no LB | WB |
| 7 | Con | 88 | M | 23 | II | 2 | neg | low | 0 | no LB | WB |
| 8 | Con | 78 | F | 34 | 0 | 1 | neg | low | 0 | No LB | WB |
| 9 | Con | 99 | F | 5 | II | 0 | neg | not | 0 | no LB | WB + MS |
| 10 | Con | 64 | M | 93 | I | 0 | neg | not | 0 | no LB | MS |
| 11 | Con | 80 | F | 25 | III | 1 | neg | low | 0 | no LB | MS |
| 12 | Con | 85 | M | 85 | I | 2 | neg | low | 0 | no LB | His |
| 13 | Con | 60 | M | 60 | 0 | 1 | neg | low | 0 | no LB | His |
| 14 | Con | 81 | F | 31 | III | 3 | neg | low | 0 | no LB | His |
| 15 | Con | 90 | F | 63 | II | 0 | neg | low | 0 | no LB | His |
| 16 | Con | 88 | M | 69 | IV | 3 | B | Inter | 0 | no LB | His |
| 17 | Con | 88 | M | 24 | II | 3 | neg | Low | 0 | no LB | His |
| 18 | DLB | 74 | M | 60 | II | 5 | neg | low | 5 | neocrtx | WB |
| 19 | DLB | 84 | M | 72 | II | 3 | neg | low | 6 | neocrtx | WB |
| 20 | DLB | 79 | M | 12 | II | 4 | neg | low | 6 | neocrtx | WB |
| 21 | DLB | 92 | F | 16 | II | 5 | neg | low | 5 | neocrtx | WB |
| 22 | DLB | 68 | M | 15 | II | 4 | A | low | 6 | neocrtx | WB |
| 22 | DLB | 73 | F | 99 | III | 4 | neg | low | 6 | neocrtx | WB + His |
| 23 | DLB | 78 | M | 8 | III | 4 | B | Inter | 6 | neocrtx | WB + MS |
| 24 | DLB | 81 | M | 26 | III | 3 | B | inter | 6 | neocrtx | WB + MS + His |
| 25 | DLB | 80 | M | 34 | III | 4 | A | low | 6 | neocrtx | MS |
| 26 | DLB | 73 | M | 47 | III | 1 | neg | low | 6 | neocrtx | His |
| 27 | DLB | 81 | M | 21 | III | 4 | A | Inter | 6 | neocrtx | His |
| 28 | DLB | 78 | F | 96 | III | N.A | A | low | 6 | neocrtx | His |
| 29 | DLB | 77 | M | 24 | III | 2 | neg | low | 6 | neocrtx | His |
| 30 | DLB | 78 | M | 83 | I | N.A | neg | low | N.A | neocrtx | His |
| 31 | DLB | 72 | M | 89 | III | 0 | neg | not | 6 | neocrtx | His |
| 32 | DLB | 91 | F | 10 | III | 4 | B | Inter | 6 | neocrtx | His |

**Supplemental Table 1. Individual Human cases.** Diagnosis (Diag), age (in years), sex, post mortem interval (PMI, in hrs) and neuropathological assessment scores for neurofibrillary tangle (NFT) Braak stage, Thal phase, Consortium to Establish a Registry for Alzheimer’s Disease (CERAD), the National Institute of Ageing – Alzheimer’s Association (NIA-AA) criteria, Lewy body (LB) Braak stage and McKeith criteria are provided. For McKeith criteria, absence of Lewy pathology (No LB), Limbic predominate and Neocortical (neoctrx) predominate are indicated. Additionally, the use of each case in western blots (WB), multi-channel fluorescence histochemistry (His) and/or mass spectrometry (MS) is also listed. N.A= not available.
