## Supplemental table 2 for "Nuclear alpha-synuclein is present in the human brain and is modified in dementia with Lewy bodies"

| Component Category | Genes | FDR |
| --- | --- | --- |
| Nuclear | | |
| Nuclear lumen | 2488 / 4917 | 2.00E-244 |
| Nucleoplasm | 2303 / 4538 | 1.10E-223 |
| Catalytic complex | 895 / 1557 | 2.90E-113 |
| Ribonucleoprotein complex | 545/785 | 4.60E-113 |
| Cell junction | 1185/ 2274 | 1.30E-111 |
| Synapse | 809/1441 | 1.70E-94 |
| Vesicle | 1911/4391 | 3.20E-90 |
| Organelle membrane | 1771/4052 | 5.80E-84 |
| Extracellular vesicle | 1130/2334 | 7.90E-80 |
| Extracellular organelle | 1130 /2336 | 1.40E-79 |
| Extracellular exosome | 1118 / 2336 | 1.00E-78 |
| Nucleolus | 601/1067 | 8.90E-70 |
| Organelle envelope | 708/1344 | 2.60E-66 |
| Envelope | 708/1344 | 2.60E-66 |
| Neuron projection | 744/1440 | 3.20E-65 |
| Mitochondrion | 885/1810 | 1.30E-63 |
| Postsynapse | 414/679 | 2.80E-60 |
| Axon | 409/895 | 8.40E-55 |
| Somatodendritic compartment | 492/895 | 2.20E-52 |
| Cell projection | 1088/2445 | 3.10E-52 |
| Transferase complex | 475/856 | 3.90E-52 |
| Anchoring junction | 494/905 | 1.90E-51 |
| Nuclear body | 505/932 | 2.30E-51 |
| Focal adhesion | 303/471 | 6.10E-51 |
| Cell-substrate junction | 306/478 | 8.70E-51 |
| Intracellular vesicle | 1196/2769 | 8.20E-50 |
| Cytoplasmic vesicle | 1193/2765 | 2.20E-49 |
| Plasma membrane bounded cell projection | 1040/2343 | 3.70E-49 |
| Dendrite | 375/649 | 1.50E-46 |
| Dendritic tree | 375/651 | 4.00E-46 |
| Cytoplasmic | | |
| Vesicle | 1911/4391 | 1.50E-254 |
| Extracellular exosome | 1189/2311 | 3.80E-219 |
| Extracellular vesicle | 1197/2334 | 3.80E-219 |
| Extracellular organelle | 1197/2336 | 6.60E-219 |
| Cell junction | 1137/2274 | 6.10E-195 |
| Synapse | 817/1441 | 4.40E-179 |
| Organelle membrane | 1575/4052 | 1.50E-140 |
| Mitochondrion | 879/1810 | 3.80E-136 |
| Cytoplasmic vesicle | 1172/2765 | 4.20E-130 |
| Intracellular vesicle | 1173/2769 | 4.80E-130 |
| Neuron projection | 708/1440 | 6.40E-111 |
| Postsynapse | 422/679 | 1.50E-107 |
| Extracellular space | 1349/3552 | 3.90E-107 |
| Cell projection | 1012/2445 | 2.90E-102 |
| Whole membrane | 881/2026 | 1.40E-101 |
| Plasma membrane bounded cell projection | 969/2343 | 3.80E-97 |
| Presynapse | 345/532 | 4.70E-95 |
| Organelle envelope | 642/1344 | 5.10E-93 |
| Envelope | 642/1344 | 5.10E-93 |
| Somatodendritic compartment | 479/895 | 1.10E-89 |
| Glutamatergic synapse | 257/353 | 1.80E-87 |
| Axon | 396/690 | 1.00E-85 |
| Neuron to neuron synapse | 262/380 | 1.10E-80 |
| Secretory vesicle | 546/1142 | 1.60E-78 |
| Mitochondrial matrix | 313/509 | 1.40E-77 |
| Dendrite | 367/649 | 7.90E-77 |
| Asymmetric synapse | 245/353 | 2.40E-76 |
| Dendritic tree | 367/651 | 2.40E-76 |
| Mitochondrial envelope | 446/868 | 7.10E-76 |
| Bounding membrane of organelle | 947/2458 | 7.50E-75 |

**Supplemental Table 2. Fraction enrichment.** Top 30 cellular component categories identified from proteomic analysis of nuclear and cytoplasmic tissue fractions as per ShinyGo database. Assigned category name, number of proteins associated genes detected out of total list are provided as well as false discovery rate (FDR).
