## Supplemental figures for "Nuclear alpha-synuclein is present in the human brain and is modified in dementia with Lewy bodies"

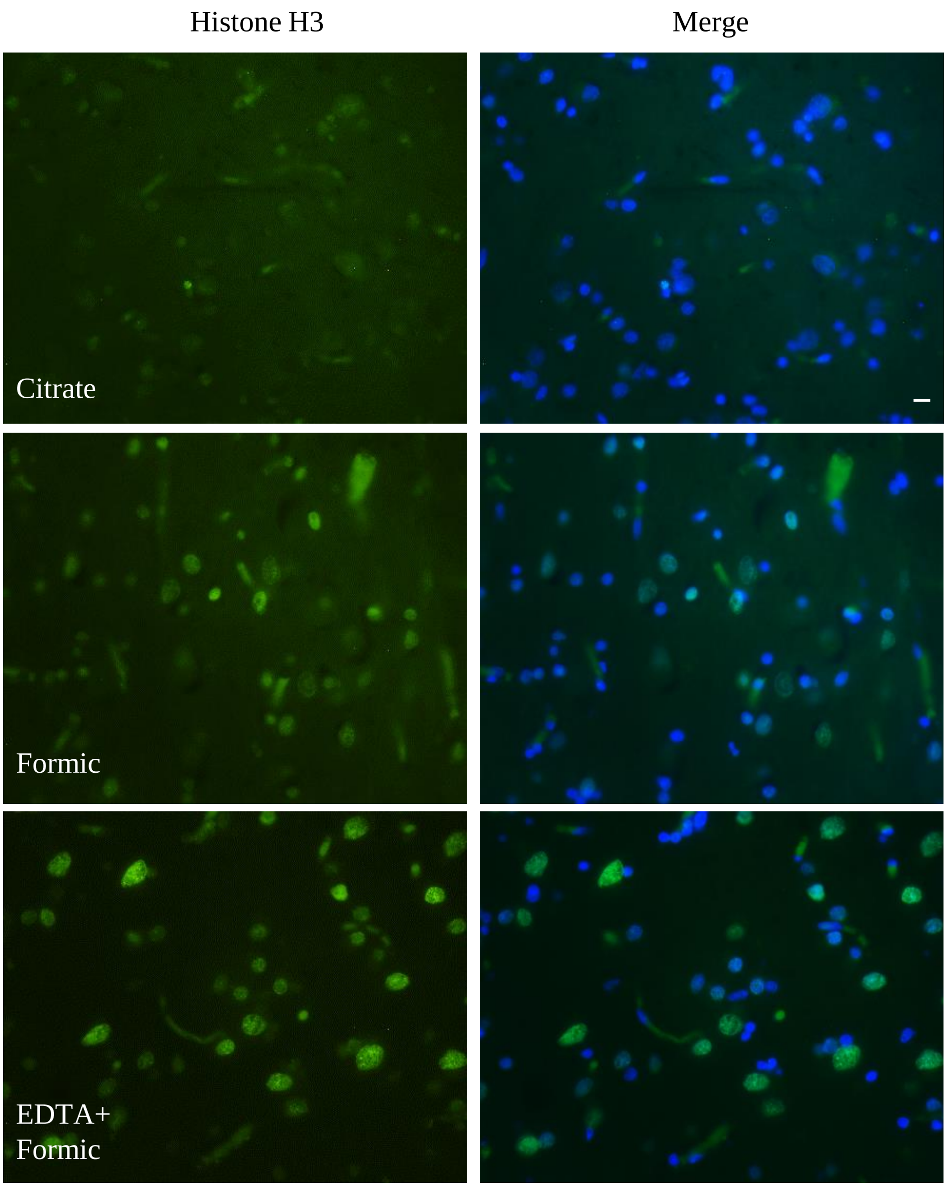


**Supplemental figure 1. Nuclear antigen accessibility following antigen retrieval methods**. Representative images captured via a 40x objective lens from control cases stained for Histone H3 with DAPI nuclear stained. Comparison of staining with citrate, formic acid and EDTA + formic acid clearly demonstrates optimum nuclear labelling following a combined treatment of EDTA and formic acid. Scale bar = 10 µm.

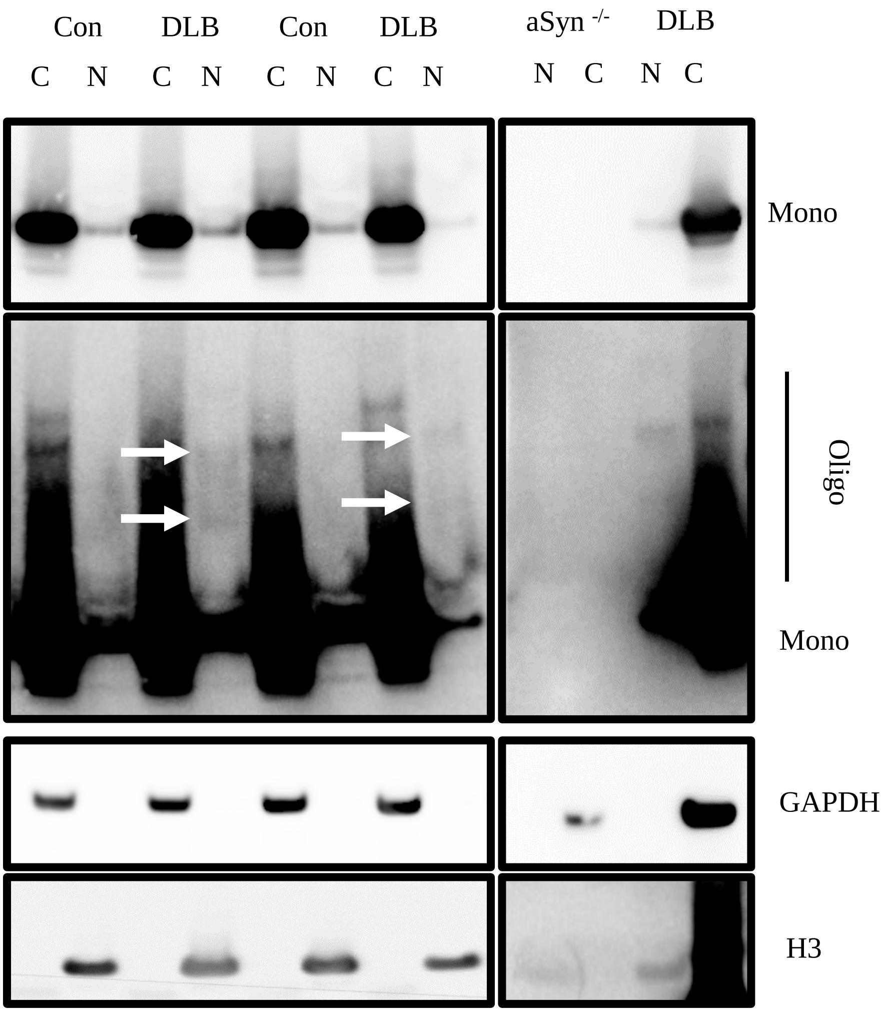


**Supplemental figure 2.Detection of nuclear aSyn via pan-aSyn antibody MJFR1**. Example western blots of pan-aSyn MJFR1 immunoreactivity of cytoplasmic (c) and nuclear (N) fractionates from control (con) and Dementia with Lewy body (DLB) cases (1.8µg/lane). Monomeric aSyn is shown under optimised exposure conditions, with large panel depicting monomeric and oligomeric aSyn species captured following overexposure of the blot. Antibody specificity was confirmed by means of similar probing of tissue fractionates generated from aSyn knockout mice (aSyn$^{-/-}$). Cytoplasmic and nuclear loading controls GAPDH and Histone H3 are also shown. Note faint appearance of high molecular weight aSyn species in the nuclear fraction only in DLB cases (arrows) only in the nuclear fraction.

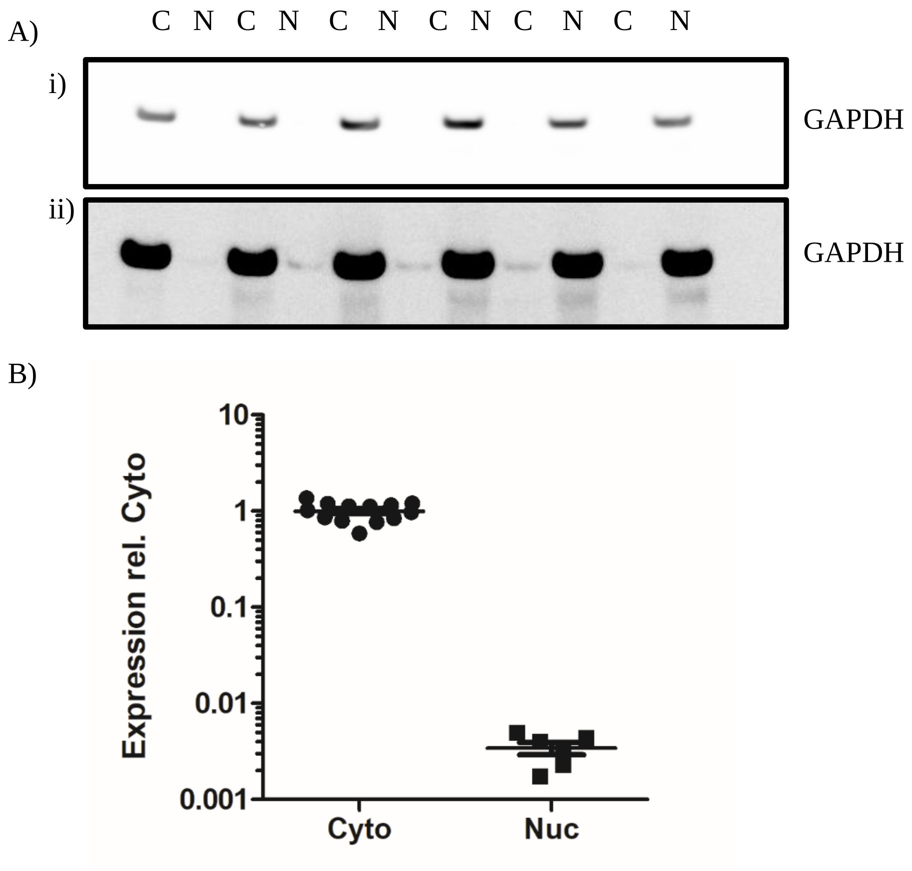


**Supplemental figure 3. Quantification of cytoplasmic proteins within nuclear fractions**. Example western blots of GAPDH immunoreactivity of cytoplasmic (c) and nuclear (N) fractionates (a) without (i) and with (ii) enhanced contrast to with enhanced contrast to allow for visualisation of GAPDH within the nuclear fraction (**A**). Comparative analysis between GAPDH immunoreactivity from cytoplasmic and those sample in which GAPDH was above detection threshold (~46%) indicated ~ a 300 fold dilution of cytoplasmic components in the nuclear fraction (**B**).

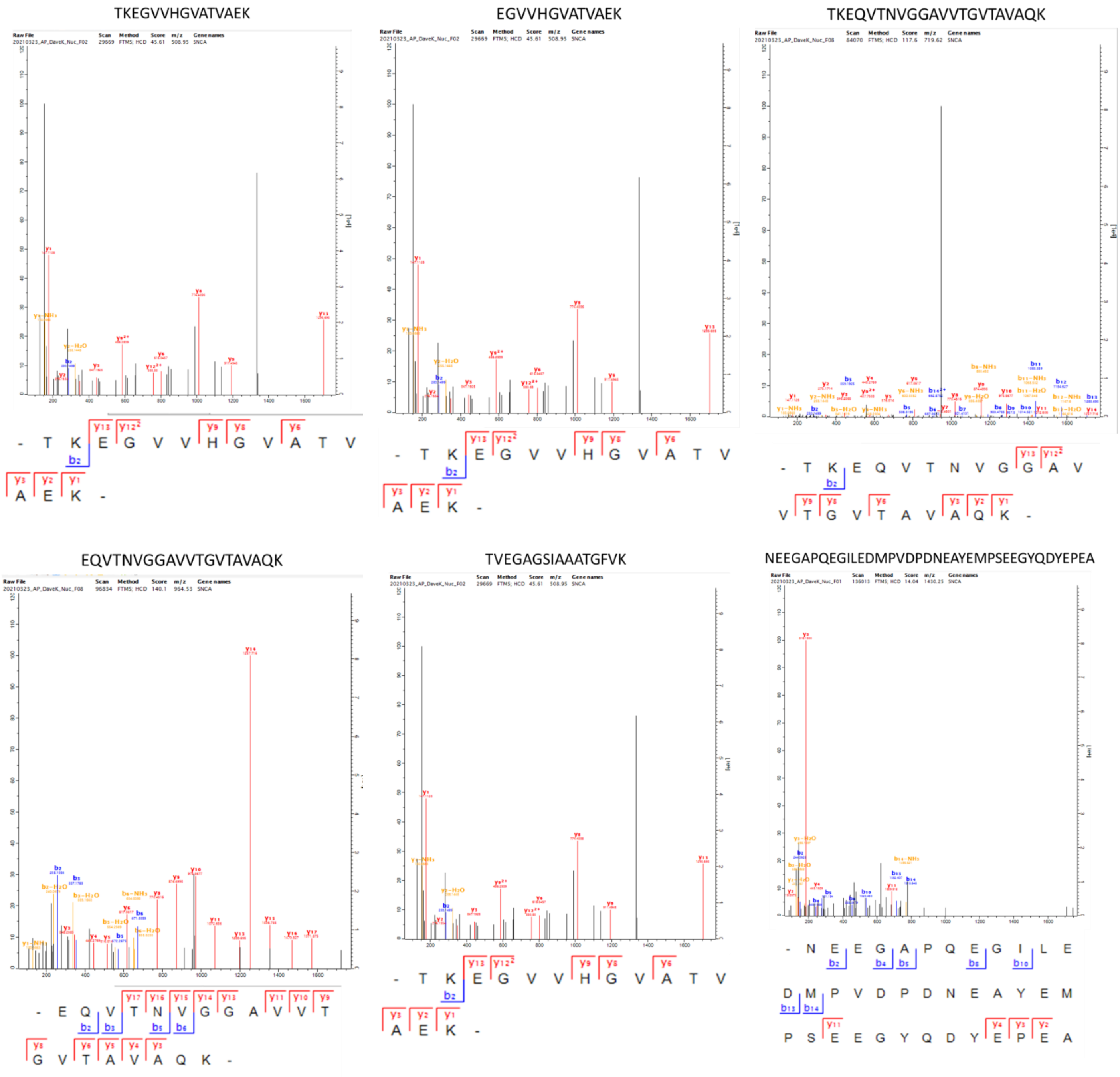


**Supplemental figure 4. Mass spectrometry detection of alpha-synuclein in the nuclear fraction**. Annotated spectra for each of the recovered peptides, b and y ions are highlighted accordingly.
